# Splicing neoepitope prediction is sensitive to methodological differences

**DOI:** 10.1101/2025.09.10.674685

**Authors:** Laurie Prélot, Julianne K. David, Andy Lin, André Kahles, Mariia Yurchikova, Reid F. Thompson, Gunnar Rätsch

## Abstract

**Motivation:** Cancer-specific neoepitopes may arise from abnormal splicing in the transcriptomic landscape (alternative splicing neoepitopes, ASNs), leading to divergent proteins with high potential immunogenicity. Predicting candidate ASNs requires a series of algorithmic and parameter choices, but no previous study has investigated the consistency and interpretability of these choices.

**Results:** We apply two ASN prediction methods with 35 matched parameter sets to generate candidate ASNs for five breast and ovarian cancer samples from The Cancer Genome Atlas (TCGA). We find that: 1) junction 9-mer peptides generated by two similarly designed pipelines differed by, on average, 68.9% and 76.6% in the BRCA and OV cohorts, respectively, and putatively cancer-specific junction 9-mers from the two pipelines diverged further, by an average of 81.5 % in OV and 84.6 % in BRCA; 2) the most lenient filters in the BRCA cohort show the highest divergence, at 97 %; 3) the rate of mass spectrometry validation of ASNs’ protein presence in cells is dominated by the size of the input space; and 4) putatively cancer-specific ASNs found by the intersection of both pipelines can be validated when accounting for false discovery with an average of 1.74 and 284.4 candidates in the BRCA and OV cohorts, respectively. Taken together, these results highlight that ASN identification is fragile and difficult to reproduce across analysis platforms, with limited cross-pipeline overlap and strong dependence on parameter choices.

**Availability and Implementation:** Python research code and scripts are available for download at https://github.com/ratschlab/projects2020_ohsu.

## Introduction

The field of cancer immunotherapy aims at activating the immune system to eliminate tumors. In particular, cancer vaccines offer the potential of personalizing a therapy to a specific patient while enabling fast and standard manufacturing. Cancer vaccine development relies on the identification of cancer-enriched or cancer-specific peptides which can trigger an immune response. These peptides are known as *neoepitopes* or *neoantigens*. Targeting cancer-mutated proteins from somatic small nucleotide variation (SNVs) [González et al., 2018, Richman et al., 2019] with cancer vaccines has had success in a clinical trial on melanoma patients [Sahin et al., 2017], but neoepitopes derived from other genomic and transcriptomic variants may be more strongly differentiated from the normal human proteome, and therefore have higher immunogenicity. Neoepitopes are often specific (a.k.a. *private*) to a single patient, although shared (a.k.a. *public*) neoepitopes derived from alternative and aberrant transcript splicing have been found to occur in various cancer types [Kahles et al., 2018, David et al., 2020]. The transcript splicing process has been observed to be strongly dysregulated in cancer [Lee and Abdel-Wahab, 2016], leading to noisy alternative splicing and aberrant splicing patterns [Escobar-Hoyos et al., 2019]. While some splicing isoforms may undergo nonsense-mediated decay (NMD) [Lejeune and Maquat, 2005], others form stable protein products [El Marabti and Younis, 2018] and are suitable candidates for vaccine integration. Nevertheless, splicing-derived neoepitopes (alternative splicing neoepitopes, ASNs) present various levels of cancer-specificity [David et al., 2020], and identification of neoepitopes requires complex biologically-informed bioinformatics pipelines.

Splicing-derived neoepitope discovery pipelines generally comprise three major steps: (1) preprocessing, which encompasses sequencing QC through alignment of the RNA-seq data, (2) generation, which includes the identification of alternative transcripts, alternative splicing events, or junctions and the translation of short peptides, and (3) filtering, which restricts the set of potential peptides to exclude those found in normal or other specified tissue samples to generate a set of candidate neoepitopes. Additional scoring often takes into account the prediction of binding to the major histocompatibility complex (MHC), immunogenicity, as well as more advanced features such as hydrophobicity, NMD, and proteasome degradation.

(1) Alignment of a sample’s RNA-seq reads and identification of reads supporting alternative splicing junctions has relied on several aligners such as STAR [Dobin et al., 2013, Li et al., 2024, Pan et al., 2023, Chai et al., 2022, Zhang et al., 2020], HISAT2 [Kim et al., 2019, Wang et al., 2019] and BWA-MEM [Vasimuddin et al., 2019, Wang et al., 2019] or the ABRA2 re-aligner [Mose et al., 2019, Chai et al., 2022]. Alignment parameters such as exclusion of multimapping reads, selection of primary or secondary alignments, and inclusion or exclusion of junctions with non-canonical splice sites will also influence the neoepitope set.

(2) The representation of splicing paths and events can involve RNA-seq supported junctions inserted into the transcripts from a given annotation file (SNAF [Li et al., 2024] via Multipath-PSI [Alt, 2018], IRIS [Pan et al., 2023] via rMATs [Shen et al., 2014], ASNEO [Zhang et al., 2020], [David et al., 2020]) or RNA-seq supported splicing paths being inserted in the transcript via a splice graph structure (Neosplice [Chai et al., 2022]). Use of a splicing graph, often constructed from the annotation and then augmented following a set of inclusion rules, can recapitulate more complex splicing patterns. These inclusion rules, such as path confidence scores used by SplAdder [Kahles et al., 2016], have strong downstream effects in a pipeline. Additional approaches include the use of transcript-centric software tools such as Cufflinks [Trapnell et al., 2010] or StringTie [Pertea et al., 2015] or the use of long read-sequencing [Li et al., 2024] to attempt to observe full-length transcript sequences directly.

Based on these representations, splicing events can be called. The splicing junctions and events are subsequently translated, either following the reading frames annotated (IRIS [Pan et al., 2023] via rMATS [Shen et al., 2014, Zhang et al., 2020, Chai et al., 2022]) or using all reading frames [Li et al., 2024] to generate small peptides of variable length (most often between 8 and 11 amino acids (AAs)).

(3) The filtering step primarily excludes candidates found in normal samples by determining their *cancer-specificity* (candidates present solely in cancer samples) or *cancer-association* (candidates present in cancer samples and with tolerated low-grade normal expression). Conversely, filtering can aim to select candidates with moderate or strong expression in the cancer sample of interest, and/or recurrence across a cancer cohort; or cancer *enrichment* (statistically significant difference between tumor and normal samples). Percent-spliced-in (PSI) [Pan et al., 2023, Zhang et al., 2020, Li et al., 2024] or normalized junction count at the RNA level [Li et al., 2024, Pan et al., 2023, Chai et al., 2022] constitute two different metrics to assess the level of splicing in cancer and normal cohorts. Based on these metrics, thresholds on expression and recurrence levels [Zhang et al., 2020, Li et al., 2024, Chai et al., 2022, Kahles et al., 2018, Pan et al., 2023], or statistical frameworks [Li et al., 2024, Pan et al., 2023] have been used to select candidates.

The choice of normal background is critical in the filtering step. The normal samples can comprise (a) RNA-seq data from a set of normal tissues processed uniformly with the foreground target data; (b) an external database of junctions (e.g. annotation [Chai et al., 2022, Zhang et al., 2020]); or (c) a set of normal peptides (e.g. UniProt [UniProt Consortium, 2023]). The size of the background as well as the type of normal tissues included is critical to the observed output. Some studies have used the GTEx [GTEx Consortium, 2013] background with about 3000 samples [Li et al., 2024, Zhang et al., 2020, Kahles et al., 2018], or about 9000 samples [Pan et al., 2023] while others have used matched tissues or cells. Several studies allow junctions defined as “cancer-specific” to be expressed in the background at levels from 1 to 4 normalized reads [Pan et al., 2023, Li et al., 2024, Chai et al., 2022, Zhang et al., 2020].

Further filtering criteria may relate to presentation of the candidate neoepitopes to T-cells and T-cell reactivity including assessment of: (a) binding to the MHC, (b) immunogenicity [Zhang et al., 2020, Li et al., 2024], (c) hydrophobicity, (d) NMD, or (d) proteasome degradation. MHC binding has been largely investigated *in-silico* [Lundegaard et al., 2008, Jurtz et al., 2017, O’Donnell et al., 2020, Shao et al., 2020]. Binding thresholds found in the literature [Li et al., 2024, Chai et al., 2022, Pan et al., 2023, Zhang et al., 2020] often comprise a *binding rank* less than 0.5-2% or an *IC50 value* of less than 50-500nM. Overall, filtering criteria are not uniformly applied and are occasionally combined into study-specific scores.

To date, published pipelines [Li et al., 2024, Pan et al., 2023, Zhang et al., 2020, Chai et al., 2022, Wang et al., 2019] have shown substantial variability in how these three steps may be implemented, with implications for downstream neoepitope prediction. Therefore, a knowledge gap remains regarding the influence of the methods and parameters used in preprocessing, generation, and filtering of the candidates.

The validation of ASN candidates relies on orthogonal observation of the candidates in -omics or experimental datasets, although datasets with RNA and protein-matched data are rare. Proteogenomics analyses [Ruggles et al., 2017, Menschaert and Fenyö, 2017, Nesvizhskii, 2010] match a database of normal peptides and neoepitopes [Zhang et al., 2020, Li et al., 2024] against experimental mass spectrometry data. Databases containing only neoepitopes (without normal peptides) have been used to increase pipelines’ sensitivity and computational speed [Li et al., 2024], but this setup does not fully control the false discovery rate (FDR) under standard target–decoy frameworks. Overall, in the context of splicing-derived neoepitopes, various search engines and target-decoy competition (TDC) approaches have been used (MaxQuant [Cox and Mann, 2008]/Andromeda [Cox et al., 2011] in SNAF [Li et al., 2024], MS-GF+ in the IRIS software tool and in a pan-cancer analysis [Pan et al., 2023, Kahles et al., 2018], Comet [Eng et al., 2013]/Percolator [Käll et al., 2007] in ASNEO [Zhang et al., 2020]), and various levels of FDR have been applied; from 1% [Zhang et al., 2020] to 5% [Li et al., 2024, Pan et al., 2023]. Notably, the FDR correction can be performed at different levels, such as the Peptide Spectrum Match (PSM) level or the peptide level [Lin et al., 2024]. Among neoepitope discovery pipelines, few are the ones which explicitly mention the type of FDR correction applied [Wang et al., 2019].

While proteomics remains one of the best tools to validate the candidates, no information is available regarding the efficacy of a candidate for tumor regression upon vaccine injection. Therefore, pipelines with heuristic steps are the best available tools to screen a large number of candidates *in-silico*. Sensitivity and specificity of pipelines for candidate detection is difficult to assess, and lack of standardization of these pipelines leads to poor understanding of the consistency of predictions across methodologies. Notably, no study to date has compared the overlap between the set of neoepitope candidates. In this study we present an exploration of the space of neoepitope candidates generated by two pipelines: David et al. [2020] and Prelot et al. [2025], including how varying parameters influence downstream predictions from mass spectrometry (MS) data. Both pipelines rely on the insertion of junctions into reference transcripts, while presenting different heuristics relative to the junction support or to the paths considered. We align our pipelines’ preprocessing steps, generate the candidates with the two pipelines, and propose a comparative analysis of the filtering parameters. Based on MS validation experiments we make recommendations to optimize the discovery of candidate neoepitopes, while limiting the probability of side effects of potential cancer vaccines.

## Results

### Systematic comparison of two ASN identification pipelines

We systematically reviewed the different design decisions of two separate ASN identification pipelines (also designated as splicing neoepitope candidate discovery pipelines), a graph-based pipeline (GP) and a junction-based pipeline (JP). The pipelines were developed using independent methods (GP: Kahles et al. [2018], Prelot et al. [2025], JP: David et al. [2020]), and here were applied to perform a series of operations conceptually as similar as possible. Despite endeavoring to align the two pipelines, we find that differences were introduced by both the junction inclusion choices (peptide generation algorithms), and the filtering heuristics (**Figure 1**, see detailed cases in Supplementary Results).

**Fig. 1.**
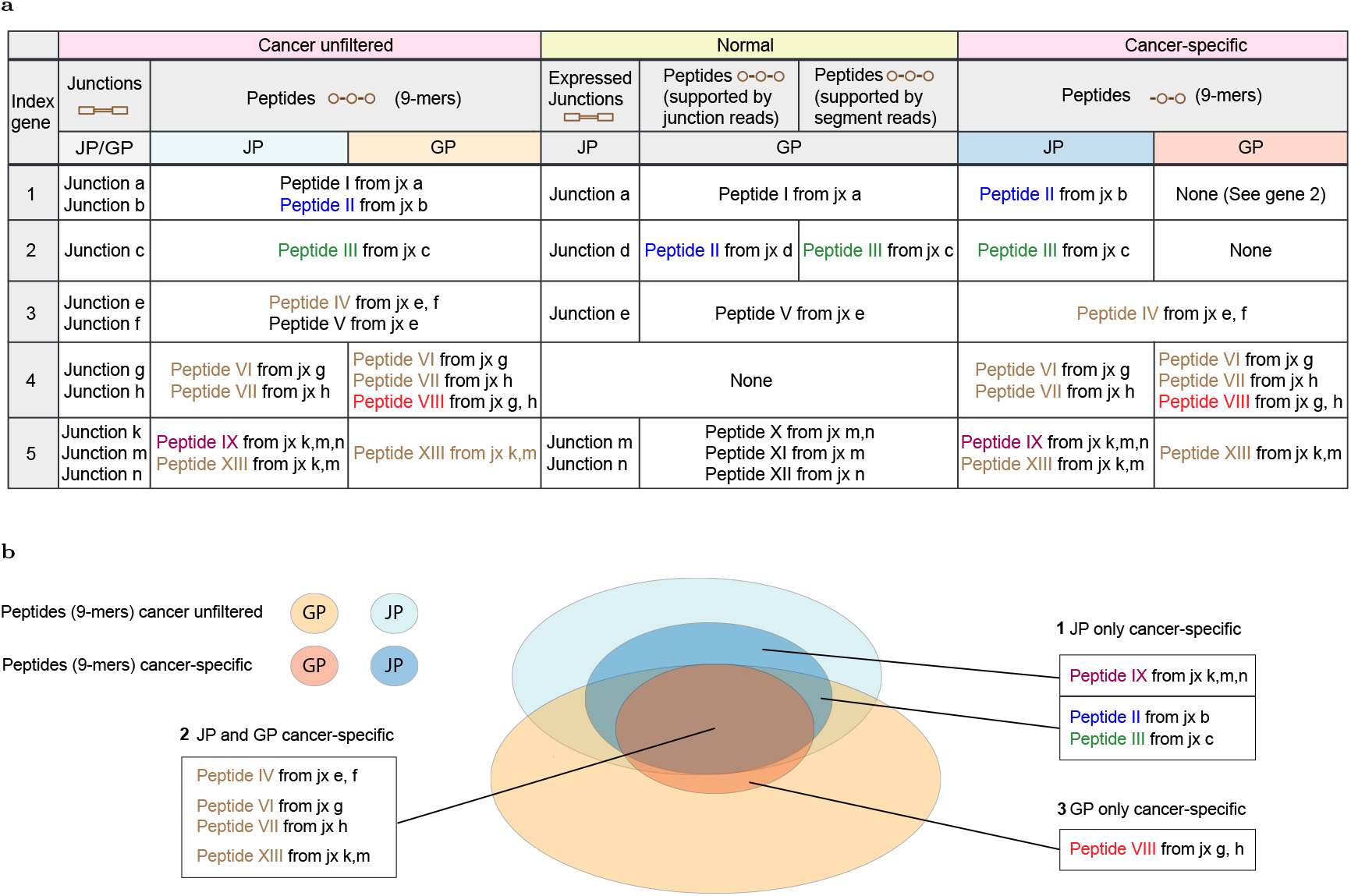
Differences between the GP and JP’s outputs illustrated on a subset of genes. (a) Table: Illustration of the emergence of discrepancies between the pipelines at different filtering and processing stages. This table shows the progression of pipeline filtering heuristics when read from left to right, showing each pipeline’s candidate peptides and how a normal-sample filter is applied to each of these. The initial set of candidates based on the RNA-seq data is “unfiltered” (left, pink) and compared to the junctions or peptides found in the normal samples (middle, green). The final set of candidates is the set of “putatively cancer-specific” candidates (right, pink). For simplicity, this table shows filtering operations on only a subset of selected candidate peptides and junctions to illustrate different outcomes or handling between and within the pipelines. Each example gene is potentially alternatively spliced, with a collection of transcripts containing exon-exon junctions. Letters are used to index junctions and correspond to different genomic positions. The translated peptides are generated from two or more exons (they contain 1 or more junctions). Roman numbers are used to index peptides and, importantly, correspond to unique peptide sequences that may arise from multiple genes or junctions across a gene or the whole genome. As the pipelines are executed independently, the set of peptides at each step can differ. Therefore, row 3 shows which pipeline a given peptide can be found in. In the case of the normal filter, this example shows that the filtering of the JP and the GP happens against any expressed junctions or peptides, respectively. Additionally, for the GP, we show in two columns the filtering against peptides supported by reads either crossing an exon-exon junction or only mapping to the body (segment) of an exon. (b) Classification of the peptides outputs from Table a based on sharedness at different steps. The Venn diagram shows the set of unfiltered (light colors) or putatively cancer-specific (dark colors) peptides for the GP and the JP. Class 1 (upper box) contains peptides present in the JP pre-filtering but not generated by the GP and classified as putatively cancer-specific by the JP. Class 1 (lower box) contains peptides present in both pipelines pre-filtering and classified as putatively cancer-specific solely by the JP. Class 2 contains peptides present in both JP and GP after filtering the candidates against the panel of normal samples following the respective pipeline strategy. Class 3 contains peptides present in the GP pre-filtering but not generated by the JP and classified as putatively cancer-specific by the GP.

The discrepancies introduced by the junction inclusion step lead to observed differences in the set of peptides prior to filtering for cancer specificity (GP-only and JP-only unfiltered sets). The sets found in this experiment are schematically represented in Figure 1 b (light colors). For instance, both pipelines generate peptides spanning one or two junctions (“Peptide IV” and “Peptide “V” in Figure 1), but only the JP can generate peptides spanning 3 junctions (“Peptide IX” is JP-specific in Figure 1). Junction insertion is performed onto a graph structure for the GP pipeline, while it is directly performed into known, annotated transcripts for the JP. Therefore, resulting peptides will diverge based on the propagation rules of the reading frame or inclusion criteria for the junctions. Junction inclusion criteria for the GP have been described in Supplementary Table 1 and [Kahles et al., 2016]. In the JP, a notable rule is that peptides from two junctions can only contain one novel junction, i.e. only one junction absent from the annotation file. An illustration is given by “Peptide VIII” in Figure 1, which does not follow the later JP-criteria and is GP-specific.

The outcome of filtering experiments, where peptides or junctions are selected based on their expression in the cancer and normal samples were then compared between the GP and JP. The filtering heuristics were closely aligned between the pipelines, but also present intrinsic differences (See Methods and Supplementary Methods 1). Their schematic view is presented in Figure 1 b (dark colors). The pipelines’ filtering outcomes present an overlap as seen with Peptides IV, VI, VII and XIII” in Figure 1, but discrepancies were observed. First, the filtering of the GP is mainly peptide-centric (Supplementary Methods), while the filtering in the JP is junction-centric. This property leads to a gene-wise candidate filtering in the JP, while the GP performs a very strict genome-wide filtering. This is illustrated by “Peptide II” in Figure 1, which is present in another gene and is filtered out by the GP. Second, the GP’s background includes both intra-exonic and junction peptides in the normal set, while the JP only includes junction peptides. This leads to a significantly larger background in the GP. Finally, the junction peptides can be quantified either on the RNA reads crossing the junction (JP, GP) or from any read overlapping any part of the gene region (GP) (Supplementary Methods). As both quantification methods are performed on the GP junction peptides, the detection sensitivity of peptides expressed in the background in the GP is increased; “Peptide III” in Figure 1 is an example.

We note that the JP pipeline has the option to filter on canonical-only splice motif junctions, which substantially reduces the number of artifactual or non-biological junctions in the output (Figure 3). However, to align the pipelines, all junctions (including those with non-canonical splice motifs) were used in downstream comparisons and proteomics validation experiments. Both pipelines filter out all peptides found in the Uniprot normal proteome.

### Generation of candidate neoepitopes by two pipelines yields different peptide spaces

We applied the two different splicing neoepitope candidate discovery pipelines to 5 ovarian (OV) and 5 breast (BRCA) cancer samples from The Cancer Genome Project (TCGA) [noa, 2022] using a unified set of matched parameters between each pipeline. In particular, we examined how varying each parameter between a low and a high value (Method Table 1), individually and combinatorially affected the resulting predictions from each pipeline in terms of both overall quantity and individual identity of candidate neoepitopes. We applied filtering in two dimensions: the first (Dimension 1) is based on positive (foreground) support for a given junction and its peptides, e.g. whether they are found in additional samples in the matched cancer cohort. The second (Dimension 2) filters negatively against junctions and peptides found in normal samples, defined as a normal background. Dimension 1 increases the likelihood that junctions identified by a pipeline are real, biological splice junctions, while dimension 2 increases the likelihood that identified junctions are truly cancer-specific and decreases the risk of off-target effects in normal tissues.

**Table 1.**
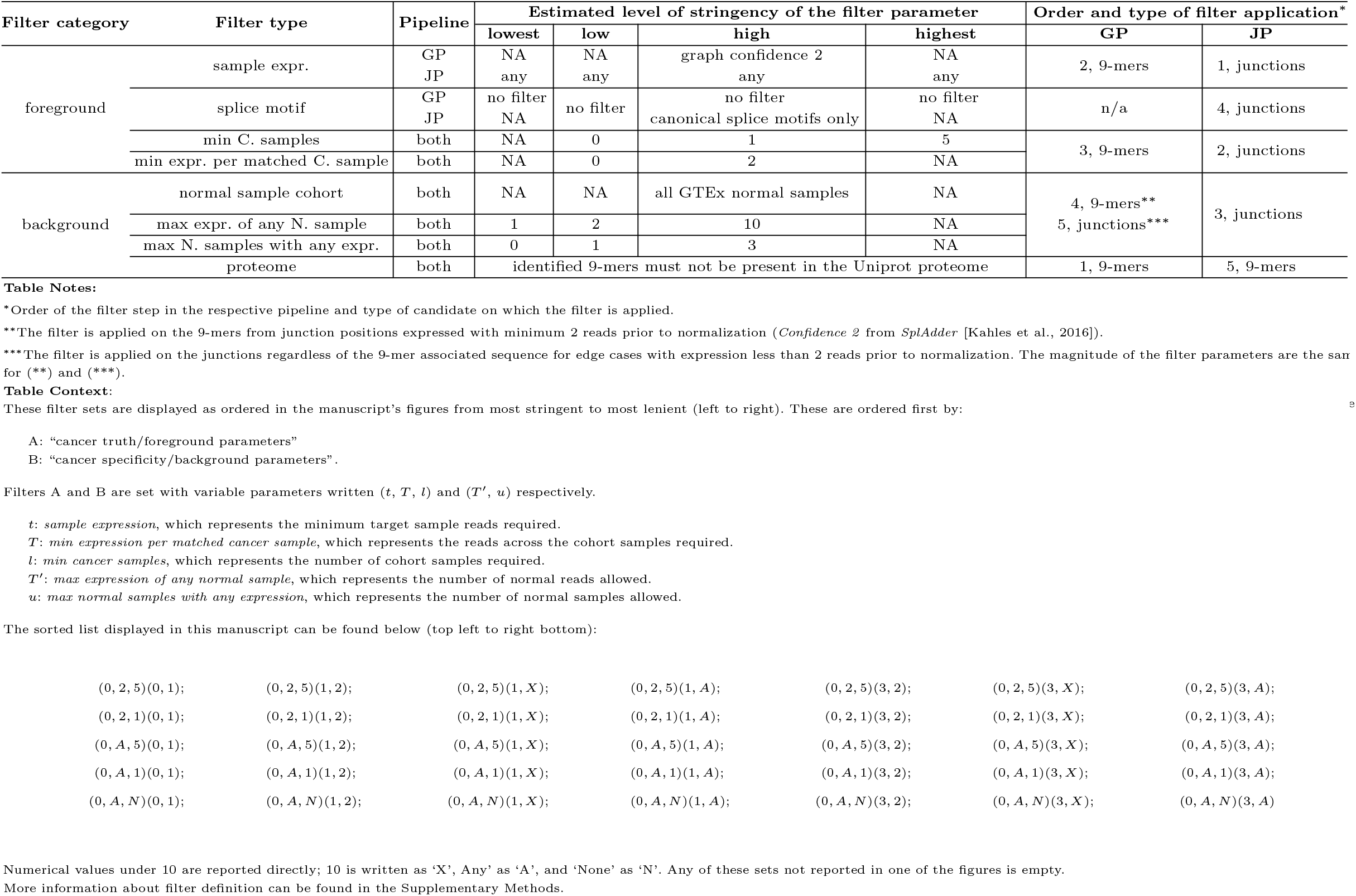
Values of filtering parameters applied on both pipelines. The foreground filters define how much support is needed to consider the junction as a biological junction present in the cancer cohort, while the background filters set the level of cancer-specificity. C.:cancer, N.:normal, expr.:expression.

We compared the number of 9-amino acid peptides (9-mers) generated by each of the pipelines and absent from UniProt [UniProt Consortium, 2023] database previous to the application of dimension 1 and 2 filtering. We found that the pipelines differ substantially in magnitude at this step with a median number of 258’169 and 633’276 of 9-mers generated from the BRCA samples for the JP and GP pipelines, respectively. The OV samples showed the same trend with a median of 807’909 9-mers in the JP pipeline and 1’196’674 in the GP pipeline (Supplementary Figure 1). The pipelines also differ in the identities of their candidate 9-mers. In BRCA samples, the overlap between the two pipelines represents on average across samples 84% of the total candidates of the JP pipeline, but only 34% of the total generated GP candidates. The discrepancy was less pronounced in the OV cohort with an overlap representing on average about 47% and 32% of the total candidates in the JP and GP pipelines respectively (Supplementary Figure 1).

### Combinatorial changes in filters can make large differences in individual pipeline output

We found that different filter parameter combinations across both filtering dimensions lead to major differences in the set of filtered 9-mer candidates. The median 9-mer candidate set size across BRCA samples analyzed varied from 19 to 7,734 candidates for the JP pipeline and from 10 to 678 candidates for the GP pipeline. A very large variance in the output set with different filtering parameter sets applied was also observed in the OV cohort with a range of 21,620 to 347,795 candidates for the JP pipeline and 18,524 to 76,226 candidates for the GP pipeline (**Figure 2** and Supplementary Figure 3). The intersection between the two pipelines was also computed for a subset of filter sets with a mean intersection across samples and filter parameter sets of 24,190.4 9-mer candidates for the OV cohort and 101.5 candidates in the BRCA cohort (Figure 2 and Supplementary Figure 3). We calculated the *dissimilarity* between the sets as the ratio of the dissymmetric difference between the JP and the GP over the size of the union. The average OV cohort dissimilarities ranged from 75% to 90% across filter sets, with higher dissimilarity generally corresponding to higher leniency in the filter parameters. In the BRCA cohort, the average dissimilarity across samples ranged from 66 % to 97 %; the lowest dissimilarity occurred with a stringent cancer-cohort filter, which required 2 reads in at least five additional BRCA samples and a lenient background filter with up to 2 reads allowed in any sample of the GTEx cohort. Overall, we observed that the JP pipeline was more lenient than the GP pipeline, generating 1.3-22.8x more 9-mer candidates for BRCA samples and 1.2-5.7x more 9-mer candidates for the OV samples.

**Fig. 2.**
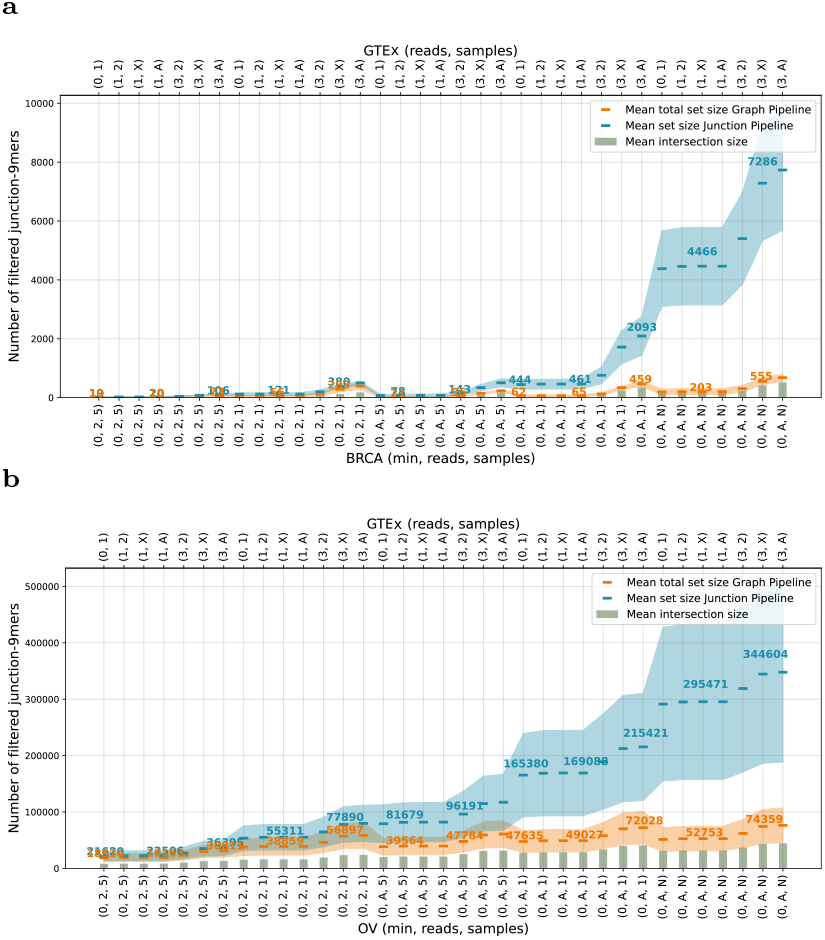
Effect of a subset of filtering parameters on the number of 9-mer candidate neoepitopes. Numbers of 9-mer candidates from the GP (orange) and JP (blue) for BRCA (a) and OV (b) samples. Within each plot, the mean value for one parameter set and one pipeline across samples is shown as a horizontal line within a ribbon representing the range of values across the 5 samples. Sage green bar plots show the mean size of the intersection between the two pipelines for each parameter set. The filter sets shown are ordered from most stringent (left) to most lenient (right). These are ordered first by cancer truth/foreground parameters (listed bottom, where the three comma-separated values are, in order, the numbers of minimum target sample reads required, reads across the cohort samples, and number of cohort samples respectively) and then by cancer specificity/background parameters (listed top, where the two comma-separated values are the number of normal reads allowed and then the number of normal samples allowed). Numerical values under 10 are reported directly; 10 is written as ‘X’, Any’ as ‘A’ and ‘None’ as ‘N’. More information in Methods Table 1.

### Filtering parameters show similar trends but different magnitudes between the two pipelines

We assessed whether conserved trends can be observed between *aligned* parameters on each pipeline. We found that for both pipelines the application of the cancer-cohort filters (Dimension 1) decreases the number of candidates up to 5 percent in the BRCA cohort and up to 30 percent in the OV cohort. The application of the GTEx normal filter leads to a significant drop of more than 80 and 40 percent in BRCA and OV cohorts, respectively (**Figure 3**, Supplementary Figures 4 and 5). The latter suggests that the GTEx-based filter (Dimension 2) is a specific dominant filter. In particular, the choice of a lenient filter, allowing the expression of 3 or more reads in the normal cohort instead of a more stringent filter of 1 read, increases the number of 9-mer candidates by at least a factor 3 in the BRCA cohort and 1.2 in the OV cohort (BRCA: 3.7x in JP and 4.6x in GP, OV: 1.2x in JP and 1.4x in GP) (Figure 2: *parameters* (3, 2) (3, X) (3, A)), demonstrating that filters, such as the GTEx-based filter, can have prevalent influences on output even when controlling for other parameter combinations.

**Fig. 3.**
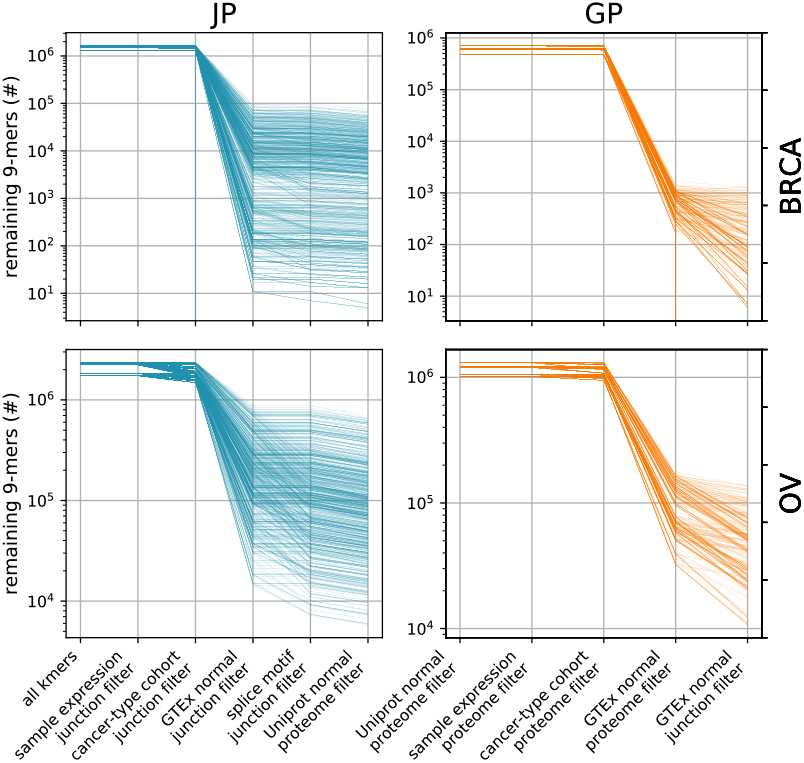
Sequential effect of the applied filters on the number of 9-mer candidate neoepitopes. (a) The absolute number (log scale) of the initial -9-mer candidates remaining in each filtering experiment at each sequential filtering step, shown for BRCA samples (top) and OV samples (bottom) for each of the JP (blue, left) and GP (orange, right) pipelines. Within each panel, each line represents the 9-mers present in one filtering experiment performed on one sample from start to finish, with 75 experiments per sample for the GP (750 total) and 180 total experiments for the JP (1800 total). This figure represents the full breadth of experiments run, including canonical splice motif and tissue-matched normal filtering for the JP, and a more broad range of normal expression levels for the GP.

### Cancer recurrence requirements affect one pipeline more strongly

Given the variation in candidate sets, we identified systematic differences related to the behavior of cancer-recurrence within the pipelines. First, the pipelines converge with stringent filters on the cancer cohort, but diverge when less (or no) recurrence is required in the associated cancer cohort (Dimension 1): an average increase in candidate count of 47x in BRCA and 5.2x in OV was observed when setting the recurrence to ‘any’, instead of 5 samples, in the JP. In the GP, decreasing the recurrence parameter within the BRCA cohort seems to have a notable, although moderate, effect (6.5x for the parameter ‘any’ relative to a recurrence of 5). This suggests that the output of the GP pipeline does satisfy recurrence in the cancer cohort (Dimension 1) prior to the combinatorial filtering step, which may be explained by intrinsic confidence criteria applied in the graph structure (Figure 2).

Conversely, the JP pipeline is very sensitive to the cancer recurrence parameter values (Dimension 1). We investigated the overlap in predicted 9-mer identity between the pipelines. We found that shared 9-mers were between 40% and 77% of the GP pipeline (40-77% for BRCA and 41-60% for OV) and 3 to 42% of the JP pipeline (3-42% for BRCA, 11-37% for OV) (Figure 2 and Supplementary Figure 3).

### Proteomics enables validation of a limited number of candidates

We attempted to validate neoepitope candidates predicted for the BRCA and OV by detecting their protein-level expression in the patients’ cells via MS data from the Clinical Proteome Tumor Analysis Consortium (CPTAC). We applied the *Subset-Neighbor-Search* [Lin et al., 2021] algorithm for proteomics because of its sensitivity to rare peptides and a stringent peptide-level FDR correction method from the *Crema* [Lin et al., 2024] tool.

In the BRCA sample set, the number of peptides validated in either pipeline is vanishingly low. The mean number of tryptic peptides spanning splice junction positions (“tryptic junction peptides”) confirmed with MS across BRCA samples varied between 0-11 and 0-1 candidates across the different experimental parameter combinations in the JP and GP, respectively (**Figure 4** A). After *in-silico* digestion of all tryptic peptides to 9-mers of 9 amino acids that span the splice junction (“junction 9-mers”), these validation numbers grew to 0-70 for the JP and 0-9 for the GP, respectively (Supplementary Figure 7 a and c).

**Fig. 4.**
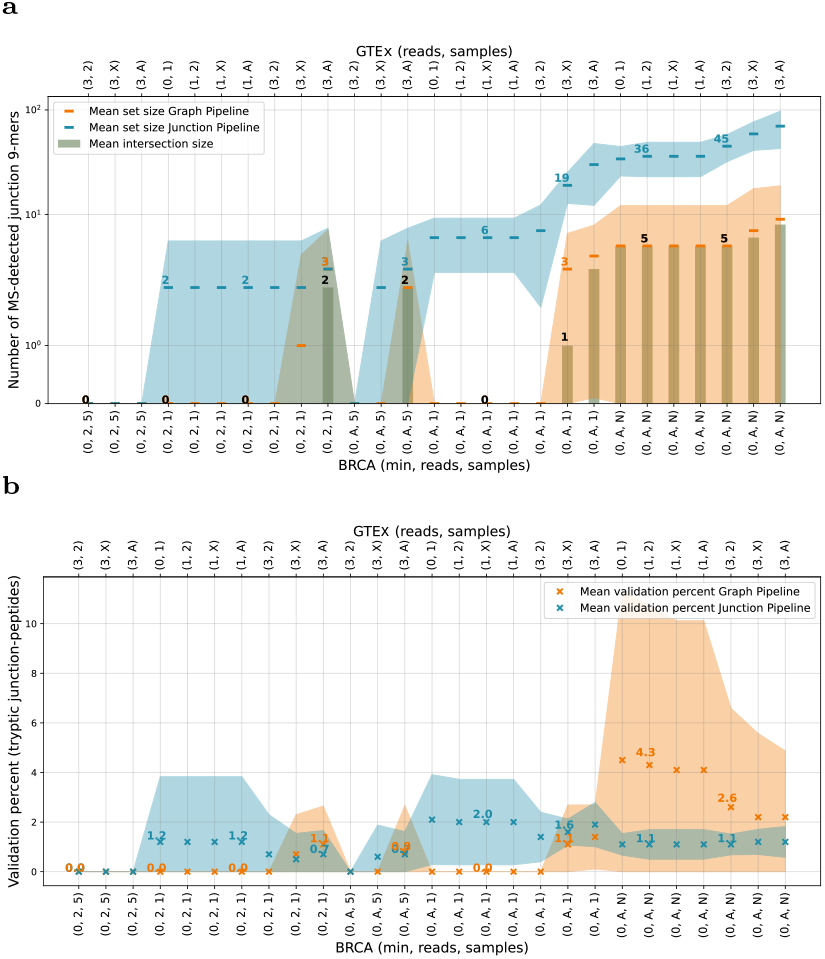
Effect of a subset of filtering parameters on MS proteomics validation for BRCA samples. Numbers of 9-mers (a) and percent of junction-overlapping tryptic peptides provided to the algorithm (b) that were validated in the GP (orange) and JP (blue) by a joint MS search across candidates from both pipelines. Subset-neighbor search was applied to an input set of junction-overlapping tryptic peptides, representing a larger set of junction-overlapping candidate 9-mers. Peptide-level FDR was calculated using *Crema*. Within each plot, the mean value for one parameter set and one pipeline across samples is shown as an “x” within a ribbon representing the range of values across the 5 samples. In subplot A, the sage green bar plots show the mean size of the intersection between the two pipelines for each parameter set. The filter sets shown are ordered from most stringent (left) to most lenient (right). These are ordered first by cancer truth/foreground parameters (listed bottom, where the three comma-separated values are, in order, the numbers of minimum target sample reads required, reads across the cohort samples, and number of cohort samples respectively) and then by cancer specificity/background parameters (listed top, where the two comma-separated values are the number of normal reads allowed and then the number of normal samples allowed). Numerical values under 10 are reported directly; 10 is written as ‘X’, Any’ as ‘A’ and ‘None’ as ‘N’. More information in Methods Table 1.

Ovarian samples had more MS-confirmed tryptic junction peptides with mean values across samples between 64 and 675 tryptic junction peptide candidates (384-4,136 junction 9-mers) across the different parameter combinations of the JP, and between 41 and 150 MS-confirmed tryptic junction peptides candidates (205-773 junction 9-mers) for the different parameter combinations of the GP. (**Figure 5** a and Supplementary Figure 8 a and c). The result in ovarian samples is consistent with the larger set of predicted filtered candidates in the OV cohort.

Substantial variability is observed between the samples within each cancer cohort, and even fewer candidates shared by both pipelines are validated with MS. The intersection of candidates confirmed with MS in both pipelines varied from 0 to 8 candidates in the BRCA cohort across filtering parameter sets (mean 1.74; for non-empty intersections the minimum was 1 candidate) and from 115 to 505 candidates in the OV cohort across filtering parameter sets (mean 284.4). For the BRCA cohort, we observed that all candidates confirmed with MS in the GP were also confirmed in the JP.

### MS peptide validation scales with cancer support and candidate counts, with confident detection lost for small input sets

In order to better understand the validation trend, we assessed the validation rate of both 9-mers (Supplementary Figures 7 d and 8 d) and tryptic junction peptides. We present primarily results for tryptic junction peptides, since these are the input seen by the MS algorithm (Figures 4 b and 5 b).

**Fig. 5.**
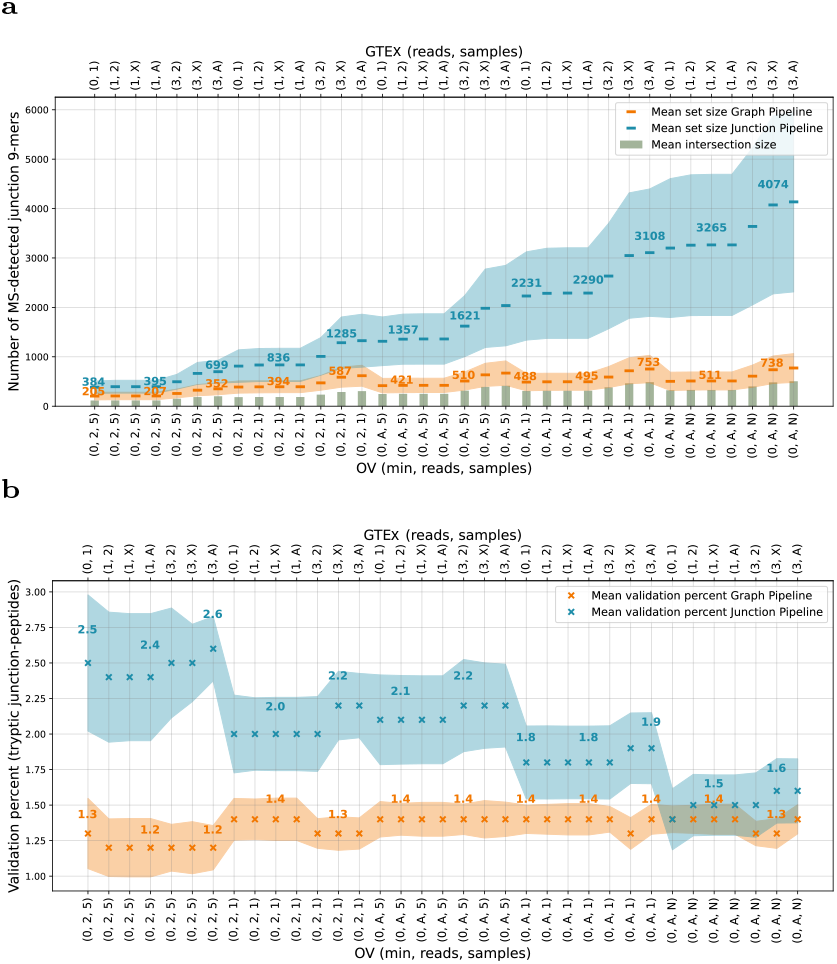
Effect of a subset of filtering parameters on MS proteomics validation for OV samples. Numbers of 9-mers (a) and percent of junction-overlapping tryptic peptides provided to the algorithm (b) that were validated in the GP (orange) and JP (blue) by a joint MS search across candidates from both pipelines. Subset-neighbor search was applied to an input set of junction-overlapping tryptic peptides, representing a larger set of junction-overlapping candidate 9-mers. Peptide-level FDR was calculated using *Crema*. Within each plot, the mean value for one parameter set and one pipeline across samples is shown as an “x” within a ribbon representing the range of values across the 5 samples. In subplot A, the sage green bar plots show the mean size of the intersection between the two pipelines for each parameter set. The filter sets shown are ordered from most stringent (left) to most lenient (right). These are ordered first by cancer truth/foreground parameters (listed bottom, where the three comma-separated values are, in order, the numbers of minimum target sample reads required, reads across the cohort samples, and number of cohort samples respectively) and then by cancer specificity/background parameters (listed top, where the two comma-separated values are the number of normal reads allowed and then the number of normal samples allowed). Numerical values under 10 are reported directly; 10 is written as ‘X’, Any’ as ‘A’ and ‘None’ as ‘N’. More information in Methods Table 1.

To better understand the validation rate, it is important to consider that the validation relies on target and decoy peptide discovery. The false discovery rate (FDR) of a set of peptide-spectrum matches (PSMs) is computed by calculating the number of decoy peptides +1 over the number of target peptides. In general, we expect that increasing the proportion of relevant peptides in the set via filtering improves sensitivity. On the other hand, low validation due to high FDR is obtained when the absolute number of targets selected by the filtering is very low. This is because numerically it is impossible to obtain a good FDR given a small number of targets. For example, given a filtering experiment that resulted in a total of 50 target peptides, the theoretically best FDR that can be obtained is 2% ([1 + 0]*/*50).

We present the results for a joint MS search across candidates from both pipelines to enable fair comparison. Overall, we observed that a joint search is very similar to pipeline-separated searches but may slightly depress validation rates when the input space is large and noisy (OV) and may support accurate validation when the number of input peptides is very low (BRCA) (Supplementary Results; see Supplementary Methods for additional details).

Across BRCA samples, the mean of the tryptic junction peptide validation rate for the JP varies from 0 and 2.1% across experimental filter sets, and for the GP varies from 0 to 4.5%. Across OV samples, this rate is between 1.4 % and 2.6% across experimental setups for the JP and between 1.2 and 1.4 % for the GP.

When using more stringent filter parameters in the JP we observed that the mean of the validation rate in the OV cohort increases; in particular, when requiring more support across external cancer cohort samples (Figure 5 b). For the BRCA cohort, the mean validation rates of the JP are universally too low to observe whether or not a similar trend exists (Figure 4 a) although some individual filters increase the validation rate. In the case of the GP pipeline, with more stringent filter parameters, the mean validation rate across experiments for the OV cohort is close to constant. This may be a result of the enrichment of the filtered 9-mer set for cancer-recurrent peptides discussed above (Figure 5 b). The GP mean of the validation rate in the BRCA cohort decreases with more stringent parameters and seems to be strongly dominated by the size of the input set: when a larger input set and no cancer-cohort sharing requirement is set, the validation is higher, likely linked to the property of the FDR calculation mentioned above for very small input sets(Figure 4 b).

Beyond the validation rates, we studied the absolute number of peptides confirmed with MS (Figure 4 a and Figure 5 a), and observed a strong trend in which the level of confirmation follows the size of the filtered input: the more candidates are fed to the MS algorithm, the higher the number of validated candidates. A slight positive effect of the cancer recurrence filter on the validation rate of the candidates is also visible.

## Discussion

This study performs a comprehensive analysis of assumptions and decisions made for cancer-specificity and cancer-association in two ASN candidate prediction pipelines. As other ASN predictions were found to be difficult to compare between pipelines with different parameter choices for the same steps [Zhang et al., 2020, Chai et al., 2022, Pan et al., 2023, Li et al., 2024], our study investigates the specific effect and possible meaning of each filtering parameter and attempts to create a set of *equivalent* or *aligned* parameters across pipelines. This study goes beyond the paradigm that pipelines generating more candidates are desirable, and attempts to better understand how the space of neoepitope candidates changes when taking different decision paths.

Our study found that different pipelines can yield very different results even with the anticipated matched parameters. In particular, differences in pipeline output at the generation stage are not the same as those that exist after applying filtering parameters. Our study also highlights that with different parameter sets used, the results from a single pipeline can show a very high degree of variation, including drastic variation in the size of the output candidate neoepitope space. Our results show how increased stringency of filtering improves the robustness of predictions, but reduces output sensitivity, in some cases resulting in no predicted candidate neoepitopes at all. Generally, the GP was found to be more strict. Heuristics driving this effect are the genome-wide filtering, the intra-exonic peptide inclusion and highly sensitive quantification for the background of the GP as noted in Results. More detailed interpretation can be found in the Supplementary Results.

Our analysis also leverages MS proteomics analysis for the validation of candidates via the *Subset-Neighbor-Search* strategy, designed specifically for rare peptides, together with the generation of *context*-inclusive multi-exon peptides for *in-silico* digestion with trypsin. Our proteomics investigation shows that support in the cancer cohort can lead to a higher likelihood of candidates’ confirmation by mass spectrometry, potentially explained by increased evidence in the patients, and by decreased competition from the normal/neighbor peptides. In our experiments, the number of validated peptides scaled with candidate-set size (Figures 4 a and 5 a), whereas validation rates changed little; (percent of junction-overlapping tryptic peptides that were validated in BRCA 0-4.5 % and 0-2.1% for GP and JP respectively, 1.2-1.4%, 1.4-2.6% for GP and JP respectively in OV, Figures 4 b and 5 b). This is consistent with target–decoy behavior at peptide-level FDR. This result suggests that the high validation rate reported in other studies [Li et al., 2024] could be driven by leniency in the cancer association or specificity thresholds.

Our work makes some parameter choices that can be placed in the context of previous studies. First, our work uses normalized junction count similar to SNAF [Li et al., 2024], IRIS [Pan et al., 2023], and Neosplice [Chai et al., 2022]. The PSI metric has also been widely used to set the filtering thresholds [Pan et al., 2023, Zhang et al., 2020, Li et al., 2024]. Our study sets thresholds on the absolute value of the normalized counts, since this is directly interpretable and transferable between cohorts; however, setting a percentage of the samples would allow working with cohorts varying in sample numbers or aberrant splicing burden. In our work, we explored various ranges of filtering against normal tissues, which recapitulate some of the choices from the literature [Li et al., 2024, Zhang et al., 2020, Chai et al., 2022, Pan et al., 2023]. Our paper includes more than 9000 GTEx samples in the panel of normal samples, similarly to IRIS [Pan et al., 2023]. In a pan-cancer analysis [Kahles et al., 2018] the cohort consists of 3,233 samples, and in the SNAF [Li et al., 2024] and ASNEO [Zhang et al., 2020] pipelines it consists of 3,000-4,000 samples. Possible decision points relative to the identity of normal samples are enrichment of the normal cohort for tissue-matched samples [Li et al., 2024], or exclusion of developmental tissues [David et al., 2020]. Exclusion of potential pre-cancerous samples from GTEx has not been done and remains difficult.

Practically, strict normal filters (with 0 reads tolerated), canonical splice-motif requirements, modest cancer-cohort support, and multi-pipeline intersection produced the most defensible sets of ASN in our study; we note that with such strict filters and very small set sizes, we found the downside of limited validation using a robust, FDR-controlled MS setup. Our study extensively describes and shares the complete protocol for analyses to identify predicted peptides in proteomics data. Immunopeptidomics datasets [Pan et al., 2023, Li et al., 2024], when available, provide potential validation of MHC-bound peptides.

Our work was limited in the parameter space explored to avoid combinatorial blow-up which would require substantial compute resources, and to simplify and control the experimental setup. This, in addition to some difficulty matching methods between two very different pipelines, led to some parameter choices that an individual pipeline would not employ when used alone, such as the JP pipeline parameters being set to include non-canonical splice motif junctions, which substantially increased the number of artifactual junctions in the downstream analyses. Some of our parameters are particularly stringent, these choices illustrate the risk of tuning parameters to desired output sizes and emphasize the need to pre-register thresholds when feasible. Our analyses differentiate between stringency of background and foreground filters, or those relative to normal samples vs. those relative to the target sample itself or its cancer cohort, or to inherent features of the target splice site. The former increases specificity against candidates that are expressed in another cell population in the body or in a specific tissue, while the latter increases the likelihood that a candidate is a true positive and enables prediction of candidates shared across many samples. Further parameters related to the MS validation could be investigated, such as thresholds for the definition of neighbor peptides, with more neighbor peptides leading to more stringent FDR control (see Supplementary Methods for further discussion of this point).

This study is also bound by inherent limitations of -omics experiments and databases. Databases are incomplete as cancer and normal space have not been fully characterized; available data is limited by existing technologies, available biological sampling, and cost burden. Our study looked at only two cancer types in TCGA and did not study additional post-filtering steps used in some splicing neoepitope pipelines, such as immunogenicity, hydrophobicity, or T-cell binding. As some of these computational methods were shown to exhibit discordances [Nguyen et al., 2023], these assessments will build an additional layer of complexity in pipeline comparison and remain to be investigated in the future. Finally, our analysis relies on the assumption that MS sensitivity can be used for candidate validation. Both experimental design [Li et al., 2021] of mass spectrometry and low correlation between protein expression and RNA-expression [Blencowe, 2017, Liu et al., 2016] challenge this assumption.

In conclusion, our study highlights substantial inconsistencies in splicing neoepitope prediction across two established pipelines and multiple tunable parameters. Our study demonstrates tunable sensitivity and specificity of predictions, though neither approach studied should be considered a *gold standard* for reliable and useful neoepitope generation. Until such an approach emerges, we advocate for a consensus approach (intersection of candidates across pipelines at the 9-mer level after identical normal filters), accepting reduced sensitivity for more robust specificity, analogous to TCGA/ICGC somatic variant consensus strategies that merge calls across multiple pipelines (e.g., requiring ≥ 2 concordant callers for SNVs) [Ellrott et al., 2018, Bailey et al., 2020, ICGC/TCGA Pan-Cancer Analysis of Whole Genomes Consortium, 2020]. Overall, ASN identification proved fragile across pipelines, with average dissimilarities of ∼ 66–97% (BRCA) and ∼ 75–90% (OV) across filter sets (**Figure 2**, underscoring limited cross-platform reproducibility and need for further research to more reliably identify ASNs.

## Methods

### Cancer Cohorts

The ovarian cohort comprised 378 OV samples from the Cancer Genome Atlas Research Network [2011]. The Breast cancer cohort comprised 1024 BRCA from the Cancer Genome Atlas Network [2012]. These two cohorts were chosen based on the availability of MS CPTAC data [Rudnick et al., 2016]. Alignment and graph generation was performed on the larger cohort. 5 samples were selected randomly for each cohort. The samples selected for the filtering experiments were: TCGA-24-1431, TCGA-24-2298, TCGA-25-1313, TCGA-25-1319, TCGA-61-2008 for the OV cohort and TCGA-A2-A0D2, TCGA-A2-A0SX, TCGA-AO-A0JM, TCGA-BH-A18V, TCGA-C8-A12P for the BRCA cohort.

### Normal Databases

#### GTEx Cohort

The GTEx background comprised 9,477 normal tissue samples downloaded from https://www.gtexportal.org/home/. The complete list of samples can be found at https://github.com/ratschlab/projects2020_ohsu?tab=readme-ov-file#ressources.

#### Uniprot

Protein sequences from human proteome were downloaded from Uniprot [UniProt Consortium, 2023]. The following proteome database was used: https://www.uniprot.org/proteomes/UP000005640.

#### Annotation

The GENCODE [Harrow et al., 2012] version 32 gtf gene annotation used for the splicing graph and the junction insertion was downloaded from https://www.gencodegenes.org/human/release_32.html.

#### Reference Genome

The reference genome GRCh38.p13.genome was downloaded from https://www.gencodegenes.org/human/release_38.html as a fasta file.

#### Alignment (GP/JP)

Alignments of the cancer and the GTEx samples were performed with STAR version 2.7.2b, the following parameters were used:

~~~
  --sjdbOverhang 100,
  --runThreadN 8,
  --outFilterMultimapScoreRange 1,
  --outFilterMultimapNmax 20,
  --outFilterMismatchNmax 10,
  --alignIntronMax 500000,
  --alignMatesGapMax 1000000,
  --sjdbScore 2,
  --alignSJDBoverhangMin 1,
  --genomeLoad NoSharedMemory,
  --readFilesCommand zcat,
  --outFilterMatchNminOverLread 0.33,
  --outFilterScoreMinOverLread 0.33,
  --outSAMstrandField intronMotif,
  --outSAMmode Full,
  --limitBAMsortRAM 7000000000,
  --outSAMattributes NH HI NM MD AS XS,
  --outSAMunmapped Within,
  --limitSjdbInsertNsj 2000000,
  --outSAMtype BAM Unsorted,
  --outSAMheaderHD @HD VN:1.4,
  --outSAMmultNmax 1
~~~

Although –outFilterMultimapNmax 20 allowed multimapping during alignment, we set –outSAMmultNmax 1 to retain only the primary alignment per read in the BAM for downstream quantification.

### Splicing graph building and propagation-aware translation (GP)

The software tool *ImmunoPepper* [Prelot et al., 2025] was used to generate candidate 9-mers sequences derived from longer bi- or 3-exons peptides (See supplementary methods). This step was performed independently for the BRCA, OV and GTEx cohort. Translation is then performed with the *ImmunoPepper* software tool [Prelot et al., 2025] (See supplementary methods). Cancer samples 9-mers overlapping splicing junctions were included in the study set, while for the GTEx cohort both junction and non-overlapping junction 9-mers were included in the normal set. Normalization of 9-mer coverage was performed as described in the supplementary methods.

### Junction extraction (JP)

All exon-exon junctions with non-zero sample (input junctions) coverage were extracted from the graphs generated with *SplAdder* version 2.4.3 [Kahles et al., 2016] for the larger cohorts of GTEx (∼ 9000 samples) and TCGA cancer samples (378 OV and 1024 BRCA samples). All junction coverages were normalized as in the supplementary methods. Splice site motifs were extracted for each junction from the reference genome with *samtools* v1.9.263. [Li et al., 2009] (See supplementary methods). The input junctions splice site coordinates were labeled based on their occurrence in the GENCODE version 32 [Harrow et al., 2012] gtf. Finally, junction-transcript pairs were translated by computationally inserting the junction coordinates into the transcript and applying coding Sequences (CDSs) from the GENCODE [Harrow et al., 2012] gtf file (See supplementary methods).

### Common Filtering Parameters (GP/JP)

For each pipeline, the parameters are:

- *sample expression* (*t*): Number of reads required as support for the 9-mer or junction, for the GP and JP respectively, in the target sample.
- *min cancer samples* (*l*): Number of non-target samples in the cancer cohort that need to reach an expression threshold (*T*, defined below).
- *min expression per matched cancer sample* (*T*): The read support threshold of the 9-mer or junction in the non-target BRCA or OV cohort samples.
- *max expression of any normal sample* (*T*^*′*^): Max number of reads that are allowed for a 9-mer or junction per sample in the normal cohort. It can also be viewed as the minimum number of reads required in any normal sample for inclusion of a 9-mer or junction in the normal background set (Normal cohort: GTEx [GTEx Consortium, 2013]).
- *max normal samples with any expression* (*u*): Max number of normal samples with expression at any read level allowed for the 9-mer or junction. It can also be viewed as the minimum recurrence at any read level that a 9-mer or junction needs to show for inclusion in the background normal cohort (Normal cohort: GTEx [GTEx Consortium, 2013]).

See more details about estimated level of stringency of the filter parameter, as well as order and scope of filter application in Method Table 1. Assuming that all filters are applied, the final set of candidates will be:

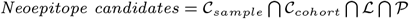

The different sets are described below:

- At the cancer level, the filtering requires a candidate 9-mer or junction to be supported in the target cancer samples with more than *t* reads. This set is named *C*_*sample*_. Besides, the 9-mer or junction is required to be expressed with *T* in at least *l* cancer samples; this set is named *C*_*cohort*_.
- At the normal level, the filtering requires the 9-mers or junctions to be excluded from the candidate set if all of the normal samples are below the expression level *T*^*′*^ (Set ℒ) or if strictly less than *u* samples are expressed with any read level (Set 𝒫).

The complete mathematical definition of the filters as well as an alternative view can be found in the supplementary methods.

### Specific Filtering steps (GP)

Prior to the parameterized filtering steps, all 9-mers found in the UniProt [UniProt Consortium, 2023] database were removed from the GP cancer set. In JP a similar step was conducted. This operation is performed to prune obvious non-neo sequences.

Parameters operate differently in JP and GP. In the JP pipeline, the cancer-specificity operates strictly at the junction coordinate level: the JP filtering is junction-centric, i.e., *candidates* = 𝒥_*S*_ (See supplementary methods). In the GP pipeline, a pre-filtering operates at the junction level for lowly expressed junctions (supplementary methods), while the main filtering criteria operate at the sequence level for 9-mers: the GP filtering is 9-mer centric, i.e., *candidates* = 𝒦 𝒥 _*S*_ .

Both methods generate candidates from canonical and non-canonical splicing.

### Dissimilarity calculation

The *dissimilarity* between the sets was defined as the ratio of the symmetric difference between the JP and the GP over the size of the union and multiplied by 100.

### Proteomics data

The proteomics data was acquired by the Pacific Northwest National Laboratory (PNNL) and Johns Hopkins University (JHU) with iTRAQ (isobaric Tags for Relative and Absolute Quantification) MS method. The data was downloaded from https://pdc.cancer.gov/pdc/ with the PDC study IDs PDC000173 (BRCA) and PDC000113 (OV).

### Proteomics steps (GP/JP)

#### General Proteomics workflow

First, tryptic peptides spanning splice junction positions (“tryptic junction peptides”) were identified. The filtered 9-mers obtained from the previous step were mapped to the longer multi-exon sequences from which they originated. From these longer multi-exon sequences, tryptic peptides were identified that contained a junction position. A tryptic peptide is defined as an amino acid sequence derived from two trypsin digestion sites or one digestion site and either the beginning or the end of a transcript. Note that the junction position could be within or between codons. In addition, any peptides with length of less than six were filtered out to reflect peptides that cannot be measured or identified using bottom-up proteomics mass spectrometers.

Next, validation of the presence of these junction sites was conducted by performing a database search on corresponding liquid-chromatography-tandem mass spectrometry (LC-MS/MS) proteomics data. In a database search, a user-defined set of peptides is searched against the experimental data. We used the subset-neighbor-search protocol [Lin et al., 2021] for our search where the target database consists of *relevant* and *neighbor* peptides. Relevant peptides are biologically relevant peptides that are expected to be present in the sample while neighbor peptides are biologically irrelevant peptides that are expected to be present in the sample and have a similar fragmentation spectrum (MS2) as a relevant peptide. For this study, we defined as relevant peptides the set of tryptic peptides defined in the previous paragraph. In addition, we defined neighbor peptides as any peptide in the human reference proteome that is similar to a relevant peptide according to the criteria in Lin et al. [2021]. Note that the search results of the fractionated samples were pooled prior to FDR control.

We used the Tide [Diament and Noble, 2011] search engine within the Crux version 4.1 toolkit [Kertesz-Farkas et al., 2023, McIlwain et al., 2014] to perform the database search. All default parameters were used except that `--precursor-window 40 --top-match 1000000000`. For the identification of neighbor peptides, we ran the function tide-index of Crux with `--mods-spec C+57.02146,K+144.102063 --nterm-peptide-mods-spec X+144.102063 --peptide-list T` followed by the peptide similarity calculation with `--min-score 0.25 --frag-bin-size 0.02 --mz-thresh 40`. More details about the neighbor peptide thresholds can be found in the supplementary methods. The false discovery rate (FDR) of the peptide detections was estimated using Crema version 0.0.9 [Lin et al., 2024] using the target-decoy competition framework [Elias and Gygi, 2007]. Specifically, peptide-level FDR was estimated using the “psm-and-peptide” methodology [Lin et al., 2022] within Crema by setting `pep fdr type=psm-peptide`. Relevant peptides passing a 5% peptide-level FDR were considered confidently detected.

#### Steps specifics to the filtering parameters comparison

Briefly, the Crux search was run with a pooling across the different parameterized filtering steps. As detailed in Supplementary Methods, pooling across filter experiments and pipelines prior to FDR control could depress apparent rates; we report them to enable fair, side-by-side comparisons. More details about the pooling can be found in the supplementary methods.

## Code availability

Python research code and scripts are available for download at https://github.com/ratschlab/projects2020_ohsu.

## Data and materials availability

Data and environments are available at https://github.com/ratschlab/projects2020ohsu?tab=readme-ov-file#ressources.

## Acknowledgments

We thank Abhinav Nellore for his involvement in the early stage of the project and his technical and strategic input and Ryan Kuck for advice on the JP codebase development. We thank Nora Toussaint for sharing her expertise on proteomics data and analyses. We give thanks to Mengyao Fan for insightful discussions about our work. We thank the Leomed 2.0 cluster from ETH Zürich for providing the compute infrastructure necessary for this work. We acknowledge the dbGAP http://www.ncbi.nlm.nih.gov/gap for providing the GTEx and TCGA data used in this manuscript with accession number phs000424 and phs000178, respectively.

