## Supplementary Material for "Splicing neoepitope prediction is sensitive to methodological differences"

### Supplementary Methods

#### Splicing graph building and propagation-aware translation (GP)

The software tool *ImmunoPepper* [Prelot et al., 2025] was used to generate candidate 9-mer sequences derived 2- or 3-exon peptides. This step was performed independently for the BRCA, OV, and GTEx cohorts. Briefly, the graph includes all the exons from the GENCODE version 32 [Harrow et al., 2012] annotation file. Then novel exons from the RNA-seq data are added iteratively to the graph. Inclusion in the graph was performed according to the read expression and anchoring criteria of *confidence level 2* of *SplAdder* version 2.4.3 [Kahles et al., 2016]. Translation is then performed with the *ImmunoPepper* [Prelot et al., 2025] software tool. For the cancer cohorts, the splicing graph was then augmented with coding Sequences (CDSs) from the GENCODE.v32 [Harrow et al., 2012] gtf file. Each CDS was propagated along the graph. For the GTEx cohort, 3 reading frames were added to each annotated exon starting from the exon start. The translation happens on a 2-exon basis following the CDSs. If the total length of two exons is less than 27 ($= 3 \cdot k$, with k=9), an additional third exon was added. The translation is interrupted if a stop codon is encountered. For each cohort, the multi-exon peptides were then split into amino acid sequences of length 9 (*9-mers)*. Any 9-mers that could be directly derived from the annotation file were removed. For the cancer samples, 9-mers overlapping splicing junctions were included in the study set, while for the GTEx cohort, both junction and non-overlapping junction 9-mers were included in the normal background set. Normalization of 9-mer coverage was performed as described below.

#### Junction extraction (JP)

The input junction splice site coordinates were labeled based on the GENCODE version 32 [Harrow et al., 2012] annotation gtf as:

- "annotated": the exact junction is present in the annotation file.
- "exon-skip annotated": both left and right splice sites are present in the annotation file, but the exact junction is not present in the annotation file (i.e. left and right splice sites do not occur in annotation on neighboring exons).
- "left- or right-splice site-only annotated": the left or right splice site is present in the annotation file, but not both.
- "unannotated": neither of the splice sites are present in the annotation file.

Genome coordinates for protein coding regions were extracted from the GENCODE.v32 [Harrow et al., 2012] gtf, and each junction was mapped to the gene IDs of any protein coding genes whose coordinate boundaries overlapped both its left and right splice sites. Each junction was further mapped to all transcript IDs for which its upstream, 5' splice site overlapped the transcript's CDS to create a set of junction-transcript ID pairs.

Junction-overlapping DNA sequence was then translated into protein sequence. For each potential junction-transcript pair, the junction was computationally inserted into the GENCODE.v32-annotated transcript as follows. If the junction's upstream splice site matched an annotated 5' splice site in the transcript, no changes were made to the upstream exon coordinate. Otherwise, the junction's upstream splice site was inserted into the upstream exon such that the upstream exon's new 3' end matched the junction's 5' end. The junction's downstream splice site was handled in one of the following ways: 1) if it already matched an annotated 3' splice site in the transcript, no changes were made to the downstream exon coordinates; 2) if it fell in the middle of an annotated exon, the upstream end of the exon was truncated so that the exon's new 5' end coincided with the 3' end of the junction; 3) if it fell between exons within the coding (CDS) boundaries of the transcript, the 5' end of the exon immediately downstream from the junction was extended to reach the junction's 3' end; or 4) if it fell beyond the end of the transcript's CDS, an artificial exon of length 150 bases was added to the end of the transcript with the new exon's 5' end matching the junction's 3' end. In every case, all exons within the junction boundaries were removed from the transcript. Each junction was considered in its matched transcripts alone; that is, even if multiple unannotated junctions were mapped to the same transcript, they were not considered in conjunction with each other.

Each *in-silico* junction-modified transcript was phased with the translation reading frame propagated from the 5' end of the original annotated transcript's coding region. On the upstream side of the junction, 150 bases were selected for translation, while on the downstream side, up to 150 bases were included according to stop codons or end-of-transcript rules. The cutoff of 150 bases was selected since tryptic peptides longer than 50 amino acids are unlikely to be detected by mass spectrometry. Note that an 300-600 base sequence could encompass any number of exons $> 2$ (joined by the target junction only), with all other junctions joining the remaining exons being fully annotated in the original transcript.

The generated sequence was then translated *in silico* with the annotated reading frame into junction-overlapping peptides of length up to 100 amino acids. The immediate peptide sequence around the junction was cut into up to 9 junction-overlapping 9-mers.

#### Relevant junction and 9-mer sets (GP/JP)

The filtering of the JP is junction-coordinate centric, while the filtering in the GP is 9-mer sequence centric. In the following paragraph, we will define junctions and 9-mer sets.

Let's define the **genome** as a continuous string of genomic positions $\mathbb{G} = g_1 g_2 \ldots g_n$, with $g_i \subset \mathbb{N}$.

Let's define **exons** as strings starting at $g_{v_{\text{start}}}$ and ending at $g_{v_{\text{end}}}$. All exons can be identified uniquely by their start and end, therefore we represent exons as pairs of genomic coordinates $v = (\text{start}, \text{end}) = (g_{v_{\text{start}}}, g_{v_{\text{end}}}) = (g_a, g_d)$ such that the pair is associated with the genomic interval $[g_{v_{\text{start}}}, g_{v_{\text{end}}}[$.

Exons are part of **transcripts**, a transcript is represented as $T_i := \{v_{i,1}, \cdots, v_{i,d_i}\}, 1 \leq i \leq m$ where $d_i$ is the total number of exons in the transcript and m is the total number of transcripts per gene. Exon pairs in a transcript are connected with exon-exon junctions. Connected exon pairs are consecutive in the transcript. It follows that a connected exon pair can be written as $(v_i, v_j) = ((g_a, g_d), (g_e, g_h))$ with $g_a < g_d < g_e < g_h$ on the positive strand. The case for the negative strand is similar.

A **junction** is defined as the pair of the last exon coordinate of the left exon and the first exon coordinate of the right exon $J_{e,b} := (g_d, g_e)$ and $g_d < g_e$.

$((g_b, g_d), (g_e, g_f))$ is a **sub-string of the exon pair** $(v_i, v_j)$. It results from the concatenation of $(g_b, g_d)$ and $(g_e, g_f)$, which are sub-strings of the exon $(g_a, g_d)$ and $(g_e, g_h)$ respectively. This means: $g_a \leq g_b < g_d < g_e \leq g_f < g_h$.

Let **alphabets** $\Sigma_{\text{RNA}}$ and $\Sigma_{\text{AA}}$ be the alphabets for RNA and protein, respectively. For a RNA string taken from $\Sigma_{\text{RNA}}$ with coordinates $((g_b, g_d), (g_e, g_f))$ and $L = (g_f - g_e) + (g_d - g_b)$ , $L \ mod(3) == 0$. The translated string in $\Sigma_{\text{AA}}$ has a length $k = L/3$ .

**Special sets**
In the following, $\Theta$ is defined as the function mapping a string given in RNA alphabet to the amino-acid alphabet.

S is a string of exactly $k$ amino acids (AAs): $S = \Theta[((g_b, g_d), (g_e, g_f))]$.

$((g_b, g_d), (g_e, g_f))$ is a sub-string of an exon pair $(v_i, v_j)$ as defined above.

We denote with $\mathcal{J}_S$ the **set of all junctions** which are contained in 9-mers of sequence exactly $S$ originating from 2-exons (in some case 3 exons). We can write:

$$\mathcal{J}_S = \{J_{e,b} | \forall((g_b, g_d), (g_e, g_f)), \Theta[((g_b, g_d), (g_e, g_f))] = S\}$$

We denote as $\mathcal{KJ}_S$ the **set of all 9-mers of sequence exactly** $S$, originating from 2 different exons, i.e. **including a junction** (in some cases originating from 3 exons). These are called *junction 9-mers*. We can write:

$$\mathcal{KJ}_S = \{((g_b, g_d), (g_e, g_f)) | \\ \forall(g_b, g_d, g_e, g_f), \Theta[((g_b, g_d), (g_e, g_f))] = S\}$$

We denote as $\mathcal{K}_S$ the **set of all 9-mers of sequence exactly** $S$, originating from genomic coordinates which **do not include a junction**. These are called *segment 9-mers*. We can write:

$$\mathcal{K}_S = \{(g_b, g_d) | \forall(g_b, g_d), \Theta[(g_b, g_d)] = S\}$$

#### Expression quantification (GP/JP)

The input junction set of JP as well as the junction and non-junction 9-mers of GP were quantified across the 378 ovarian, 1,024 breast cancer samples, and 9477 GTEx samples. Expression values were associated with peptide sequences or coordinates as follows:

(1) The **"junction expression"** or **"junction read count"** is defined as the number of RNA-seq reads that overlap a junction. These were quantified directly from the STAR junction coverages. The normalized junction expression is the read count divided by the *library size* in the sample multiplied by 400,000. This operation brings back the normalized counts to an interpretable value; the multiplier 400,000 was chosen based on empirical tests. For each sample, a *library size* normalization factor is defined as the 75th percentile expression value across all coding genes. Let $\Phi$ be the mapping associating its expression $q \in \mathbb{R}$ to the junction $J_{e,b}$.

$$\Phi(J_{e,b}) = \Phi((g_d, g_e)) = q$$

The "junction expression" value is used in the JP. By extension, the "junction expression" can also be associated to the 9-mer $((g_b, g_d), (g_e, g_f)) \in \mathcal{KJ}_S$ containing a junction $J_{e,b}$:

$$\Phi(((g_b, g_d), (g_e, g_f))) = \Phi(J_{e,b}) = \Phi((g_d, g_e)) = q$$

The "junction expression" value applied to 9-mers is used to quantify the junction-overlapping 9-mers of the GP.

(2) Each exon pair $(v_i, v_j)$ also has a **"segment expression"** or **"segment read count"**. It represents the average expression of the corresponding genomic sub-sequence. We follow the definition of segments in [Kahles et al., 2016]: An exon $v_i$ is composed of segments $s_{i,q}$ through $s_{i,r}$, if $v_i = s_{i,q} \circ_{i,r}$. Hereby, $\circ$ denotes the concatenation of segment positions. The expression counts and segment length for segment $s_{i,q}$ are $EC_{i,q}$ and $SL_{i,q}$. The length for the exons $v_i$ and $v_j$ are $VL_i$ and $VL_j$, with $VL_i := \sum_{k=q_i}^{r_i} SL_{i,k}$. The segment expression for a translated exon pair $(v_i, v_j)$ is defined as:

$$SE_{ij} := \frac{1}{VL_i + VL_j} (\sum_{k=q_i}^{r_i} SL_{i,k} EC_{i,k} + \sum_{k=q_j}^{r_j} SL_{j,k} EC_{j,k})$$

Each *segment* in the splicing graph stores an expression value extracted from the initial STAR alignment.

Note: The formula above was used to compute the gene expression of all coding genes prior to *library size* computation. The *library size* is introduced above.

By extension, for a 9-mer $(g_b, g_d) \in \mathcal{K}_S$, the expression will be computed similarly, except that if only a fraction of a segment $s_{i,k}$ is included in the 9-mer, $SL_{i,k}$ will be replaced by $SL'_{i,k}$ , the truncated segment length. $VL_i$ becomes $VL'_i := \sum_{k=q_i}^{r_i} SL'_{i,k}$ and the expression counts value $EC_{i,k}$ will remain the same.

Let $\Psi$ be the mapping associating a 9-mer to its "segment expression" $q \in \mathbb{R}$.

$$\Psi((g_b, g_d)) = q$$

The "segment expression" value applied to the 9-mers is used to quantify non-junction-overlapping 9-mers of the GP.

#### Common filtering parameters (GP/JP)

**Expression across cohorts**
The filtering includes expression filters at the sample and the cohort level. At the sample level, let's define the following **expression values**:
$E^{\gamma}$: "junction expression" value in one sample.
$E^{\sigma}$: "segment expression" value in one sample.

At the cohort level, let's define the following **expression vectors**:
$E^{\gamma}_C$: "junction expression" vector in cancer cohort.
$E^{\gamma}_N$: "junction expression" vector in normal cohort.
$E^{\sigma}_C$: "segment expression" vector in cancer cohort.
$E^{\sigma}_N$: "segment expression" vector in normal cohort.

JP and GPs first implement basic operations on expression vectors. Let's consider a normalized expression metric $q \in \mathbb{R}$ over a cohort $G$ of $n$ samples. It can be junction or segment expression.

Let $E_{\lambda,G} \in \mathbb{R}^n$ , $E_{\lambda,G} = [e_1 \ e_i \ \cdots \ e_n]$ be defined as the **normalized expression vector of** $\lambda$**, either a junction or a 9-mer candidate**, taken across $n$ samples for some theoretical cohort $G$ independently of whether the cohort is cancer or normal. $\mathbb{1}$ is the indicator vector.

We denote as $E_{\lambda,G}(T,l)^+$ the expression vectors associated with the candidates $\lambda$ where $l$ or more samples have a normalized expression greater than a user-defined threshold $T$.

$$E_{\lambda,G}(T,l)^+ := \{E_{\lambda,G} \in \mathbb{R}^n | \Sigma_{i=1}^n \mathbb{1}_{e_i > T} \geq l\}$$

Similarly, $E_{\lambda,G}(T,l)^*$ the expression vectors associated with the candidates $\lambda$ where $l$ or more samples have a normalized expression greater **or equal** to $T$:

$$E_{\lambda,G}(T,l)^* := \{E_{\lambda,G} \in \mathbb{R}^n | \Sigma_{i=1}^n \mathbb{1}_{e_i \geq T} \geq l\}$$

The special case where any sample needs to pass a user-defined threshold can be written as:

$$E_{\lambda,G}(T,1)^* := \{E_{\lambda,G} \in \mathbb{R}^n | \Sigma_{i=1}^n \mathbb{1}_{e_i \geq T} \geq 1\}$$

The special case where $l$ or more samples are expressed with any read count is written as:

$$E_{\lambda,G}(0,l)^+ := \{E_{\lambda,G} \in \mathbb{R}^n | \Sigma_{i=1}^n \mathbb{1}_{e_i > 0} \geq l\}$$

Conversely, if strictly less than $l$ samples have an expression greater than $T$, we introduce the (strict) maximal recurrence $l$ observed for an expression level of $T$:

$$\overline{E_{\lambda,G}(T,l)^+} := \{E_{\lambda,G} \in \mathbb{R}^n | \Sigma_{i=1}^n \mathbb{1}_{e_i > T} < l\}$$

The special case of (strict) maximal recurrence $l$ at read level 0 will be written as:

$$\overline{E_{\lambda,G}(0,l)^+} := \{E_{\lambda,G} \in \mathbb{R}^n | \Sigma_{i=1}^n \mathbb{1}_{e_i > 0} < l\}$$

When the maximum is set over the expression value instead of the recurrence value, the (strict) maximal expression in the cohort can be written as:

$$E_{\lambda,G}(T)^{\vee} = \{E_{\lambda,G} \in \mathbb{R}^n | \forall j \in \{1...n\}, e_j < T\}$$

**Filtering setups**

For the GP and JP experiments, we set the filtering parameters as described in the methods.

**Cancer sample and cancer cohort filtering**

The goal of the filtering is to create a "short-list" of candidates for a cancer sample of interest. This sample of interest is also referred there as the "target cancer sample". The candidates $\lambda$ in the JP are junctions ($\lambda \in \mathcal{J}_S$), while the candidates $\lambda$ in the GP are 9-mers overlapping a junction position ($\lambda \in \mathcal{KJ}_S$). The parameters set at this step are:

- $t$: *sample expression*
- $T$: *min expression per matched cancer sample*
- $l$: *min cancer samples*

The following filtering steps were implemented at the cancer level:
The first filter selects candidates present in the target sample with more than $t$ reads. This set of candidates is $\mathcal{C}_{sample}$, with $\mathcal{C}_{sample} := \{\text{candidates } | E^{\gamma} > \text{t}\}$.

The second filter selects targets for which an expression greater or equal to $T$ in $l$ or more samples of the cancer cohort. This set of candidates is $\mathcal{C}_{cohort}$, with $\mathcal{C}_{cohort} := \{\text{candidates } | E^{\gamma}_{\lambda,C}(T,l)^*\}$.

When both cancer-level filters are applied, the set of selected candidates is the intersection of the candidates passing the first filter (cancer sample-level) and the candidates passing the second filter (cancer cohort-level): $\mathcal{C} := \mathcal{C}_{sample} \bigcap \mathcal{C}_{cohort}$.

**Normal (GTEx) cohort filtering**

The goal is to continue filtering the target sample candidate "short-list". The normal-level filtering aims at excluding candidates present with "sufficient" support in the normal cohort. As above, candidates in JP are junctions, while candidates in GP are 9-mers overlapping a junction position. The parameters set at this step are:

- $T'$: *max expression of any normal sample*
- $u$: *max normal samples with any expression*

*In the paragraphs below, we focus on the **normal cohort side** and define what "sufficient" support in normal samples means.*

The first filter selects normal junctions or normal 9-mers *confidently expressed* in the normal cohort. These candidates have an expression greater or equal to $T'$ in at least one normal sample. This set of candidates is $\mathcal{T}$, with $\mathcal{T} := \{\text{GTEx junctions or 9-mers } | E_{\lambda,GTEx}(T',1)^*\}$.

In particular:

- for the GP, the candidates $\lambda$ are 9-mers that satisfy the criteria $\{\{\lambda \in \mathcal{KJ}_S | E^{\gamma}_{\lambda,GTEx}(T',1)^*\} \bigcup \{\lambda \in \mathcal{K}_S | E^{\sigma}_{\lambda,GTEx}(T',1)^*\}\}$
- for the JP, the candidates $\lambda$ are junctions that satisfy the criteria $\{\lambda \in \mathcal{J}_S | E^{\gamma}_{\lambda,GTEx}(T',1)^*\}$.

The second filter selects the normal junctions or normal 9-mers *recurrent* in the normal cohort. These candidates are found expressed in $u$ or more normal samples regardless of the expression value. This set of candidates is $\mathcal{R}$, with $\mathcal{R} := \{\text{GTEx junctions or 9-mers } | E_{\lambda,GTEx}(0,v)^+\}$.

In particular:

- for the GP, the candidates $\lambda$ are 9-mers that satisfy the criteria $\{\{\lambda \in \mathcal{KJ}_S | E^{\gamma}_{\lambda,GTEx}(0,v)^+\} \bigcup \{\lambda \in \mathcal{K}_S | E^{\sigma}_{\lambda,GTEx}(0,v)^+\}\}$
- for the JP, the candidates $\lambda$ are junctions that satisfy the criteria $\{\lambda \in \mathcal{J}_S | E^{\gamma}_{\lambda,GTEx}(0,v)^+\}$.

When both normal-level filters are applied, candidates present with "sufficient" support in the normal cohort are removed from our previously defined cancer-level filtered candidate set $\mathcal{C}$. The subtraction operation $\mathcal{C} \backslash \{\mathcal{T} \bigcup \mathcal{R}\}$ is performed to remove the normal background. In some cases, only the first filter (on normal expression level) or the second filter (on normal recurrence) are applied. There subtraction operations will be $\mathcal{C} \backslash \mathcal{T}$ or $\mathcal{C} \backslash \mathcal{R}$, respectively.

Another way of writing the normal filter is to focus on the foreground view and see the normal parameters $T'$ and $u$ as maximum levels ”tolerated” in the normal cohort.

*In the paragraphs below, we focus on the* ***cancer side view*** *and define what the maximum levels ”tolerated” in the normal cohort means.*

The first filter selects candidates from the cancer target sample that are *lowly expressed* in the normal set. The candidates' expression is strictly below the maximal value $T'$ in the normal cohort. This set of candidates is $\mathcal{L}$, with $\{\text{candidates } | E_{\lambda,GTEx}(T\prime)^{\text{v}}\}$.

The second filter selects candidates from the cancer target sample that are *lowly recurrent* in the normal set. The candidates' recurrence is strictly below the maximal value $u$ in the normal cohort. The recurrence of a candidate is calculated as the number of samples with any number of normalized reads. This set of candidates is $\mathcal{P}$, with $\mathcal{P} := \{\text{candidates } | \overline{E_{\lambda,GTEx}(0, v)^{+}}\}$.

When both normal-level filters are applied, our previously defined cancer-level filtered candidate set $\mathcal{C}$ is intersected with *lowly expressed* and *lowly recurrent* candidates: $\mathcal{C} \bigcap \mathcal{L} \bigcap \mathcal{P}$. In some cases, only the first filter (on normal expression level) or the second filter (on normal recurrence) are applied. There intersection operations will be $\mathcal{C} \bigcap \mathcal{L}$ or $\mathcal{C} \bigcap \mathcal{P}$, respectively.

In the GP, a correction was applied. As the splicing graph generated with SplAdder version 2.4.3 [Kahles et al., 2016] (confidence parameter: 2) excludes junctions expressed with less than 2 reads, the expression and recurrence pattern of those junctions in the GTEx cohort were extracted from the STAR [Dobin et al., 2013] alignment output files. Candidate junction 9-mers containing a junction present in the GTEx set were filtered out if they met the ”normal cohort filtering thresholds” described above.

#### Proteomics steps (GP/JP)

The goal of this study is to best assess the effect of various filtering parameters on the landscape of neoepitope candidates. To best compare the MS validation rates between experiments, we performed the pooling of tryptic junction peptides with two different strategies:

1. *Joint*: Peptide sets of all experiments were pooled across both pipelines JP and GP. This allowed us to analyze the validation rate of each of the pipelines in a comparable way.
2. *Separate*: The results of all experiments were pooled while keeping the peptide sets for the JP and the GP separate. This allowed us to quantify the validations as they would be for two independent studies.

Note that the choice of pooling across experiment parameters might affect FDR control. Nevertheless, we believe that this is a reasonable choice due to the small size of some filtered peptide sets. Search with the Crux version 4.1 [Kertesz-Farkas et al., 2023, McIlwain et al., 2014] search engine was performed on the sets for each strategy; (1) and (2). This included the neighbor peptides. Next, the Crux search results were concatenated across fractions and the target peptides initially from the JP and the GP were separated. The target peptides were re-ranked based on the cross-correlation score ('xcorr score' column) within each of the pipelines. The FDR was estimated with Crema version 0.0.9 [Lin et al., 2024] in *peptide-PSM* mode on the JP and GP set respectively.

The *joint* strategy yielded the same trends as the *separate* strategy. Overall, we observed that the *joint* search reduced the validation rate in OV for the JP, where the input space was very large; we hypothesize that this (1) creates more unrelated peptides that can be matched by chance and (2) generates more matches to homologous peptides that need to be controlled for (neighbor peptides). In the BRCA cohort, we observed that the *joint* search slightly increased the validation rate compared to the *separate* strategy. This could be explained by the fact that when very few relevant peptides are present, the false discovery rate will be high. As we explain in the main text, it is numerically impossible to obtain a good FDR given a small number of targets. Therefore, adding a few relevant peptides in the search space increases discovery in this case (Supplementary Figures 10 and 9).

This study aimed to include as many relevant parameter thresholds as possible, but did not empirically investigate the influence of parameters related to MS validation. Nevertheless, the neighbor peptides threshold parameters are expected to influence the FDR control. Indeed, the definition of neighbor peptides is based on (1) the proportion of $b$- and $y$-ions shared two peptides being greater than a threshold $t_i$ (2) the difference in the associated peptide masses $m_1$ and $m_2$, specified in units of ppm, being less or equal to a specified mass tolerance $t_m$. In this work, we set $t_m$ equal to twice the precursor mass tolerance of 40 ppm, and we set $t_i = 0.25$ following the settings used by Lin et al. [2021]. From the definitions, we can predict that the number of neighbor peptides will increase with (2) and decrease with (1) [Lin et al., 2021]. With more peptides identified as neighbors it becomes harder to reach a given FDR, so fewer peptide identifications will be made.

### Supplementary Results

#### Intersection between the GP and JP's outputs illustrated on a subset of genes

Peptides from the GP and the JP are derived from the same alignments and can differ due to 1) the implementation of a graph junction-insertion algorithm followed by propagation of the reading frame on the graph for the GP vs. 2) the junction insertion directly into the reference annotation transcript and translation for the JP. The intersection of the candidate sets at the generation stage is displayed in Supplementary Figures 1, 2 and schematically in Main Figure 1 b. After filtering, the peptides can be either retained by both pipelines (Class 2 in Main Figure 1 b) or retained by a single pipeline (Class 1 and 3 in Main Figure 1 b). In the latter case, the peptides may have been generated in a pipeline-specific way (Class 1 - upper box - and 3 in Main Figure 1 b), or may have been generated by both pipelines but subject to pipeline-specific filtering (Class 1 - lower box in Main Figure 1 b). The peptides in class 1 (lower box in Main Figure 1 b) are examples of peptides that are retained by the JP pipeline only.

#### Individual cases between the GP and JP's outputs illustrated on a subset of genes

In Main Figure 1 a, we show potential convergence and differences between the pipelines. As a baseline, peptide “I from junction a” is filtered out in both the GP and JP because the peptide (or junction, respectively) is expressed with any read in at least one normal sample. More complex expression thresholds are omitted for simplicity. We thereby illustrate peptide-centric filtering in the GP and junction-centric filtering in the JP (See  and supplementary methods). In Main Figure 1 a, we observe that "peptide II from junction b" in gene 1 is filtered against "peptide II from junction b" in gene 2 due to matching peptide sequence (Main Figure 1 a). This shows the genome-wide and gene-wise filtering heuristics of the GP and JP, respectively.

In the JP all normal background peptides are derived from junctions while in the GP intra-exonic peptides are included in the background set (Not shown in Main Figure 1 a).

Furthermore, "peptide III from junction c" (Main Figure 1 a) illustrates another discrepancy. When quantified by reads on the left and right side, the peptide is found to be present with enough support in the normal background set and is excluded from the GP while included in the JP because no reads are found that explicitly support the junction in the normal set.

Next, we focus on the generation of peptides from 2 junctions and show that "peptide IV from junction e and f" is retained in the set of candidates for both pipelines because solely the single-junction peptide "peptide V from junction e" (Main Figure 1 a) has been found in at least one normal sample. A particular case of a peptide from 2 junctions is: "peptide VIII from junctions g, h" (Main Figure 1 a), which is not included in the set of generated candidates for the JP because the method only considers peptides with a single potentially novel junction, with all other junctions in the peptide found in the GENCODE.v32 annotation (Harrow et al. [2012]). The GP pipeline, which benefits from cross-sample transcript sharing will include the peptide in the generated set. Finally, we focus on peptides from 3 junctions with "peptide IX from junctions k, m, n" (Main Figure 1 a), a peptide specifically translated by the JP, as the GP does not, at the moment, support concatenation of more than 3 exons.

#### Summary of differences in GP and JP's outputs illustrated on a subset of genes

Explaining the discrepancy between the pipelines is a complex task, however, presenting features that lead to differences between the JP and the GP output sets can help. On the one hand, the size of the GP output is significantly smaller relative to the JP set due to the strong effect of the genome-wide filtering in the GP. Moreover, the inclusion of non-junction background peptides beyond those present in Uniprot leads to the removal of a large number of candidates in the GP. This feature is coupled with the highly sensitive quantification of junction peptides in the GP background, which further decreases potential false positive inclusion. The JP output set is also larger than the GP output set because of the inclusion of peptides from 3 junctions by the JP pipeline. On the other hand, the size of the GP is increased relative to the JP due to the complex propagation algorithm on the graph. Reinforcing this effect, there exist potential peptide candidates in the GP that arise from two potentially novel junctions, which are not included in the JP.

#### Supplementary Figures

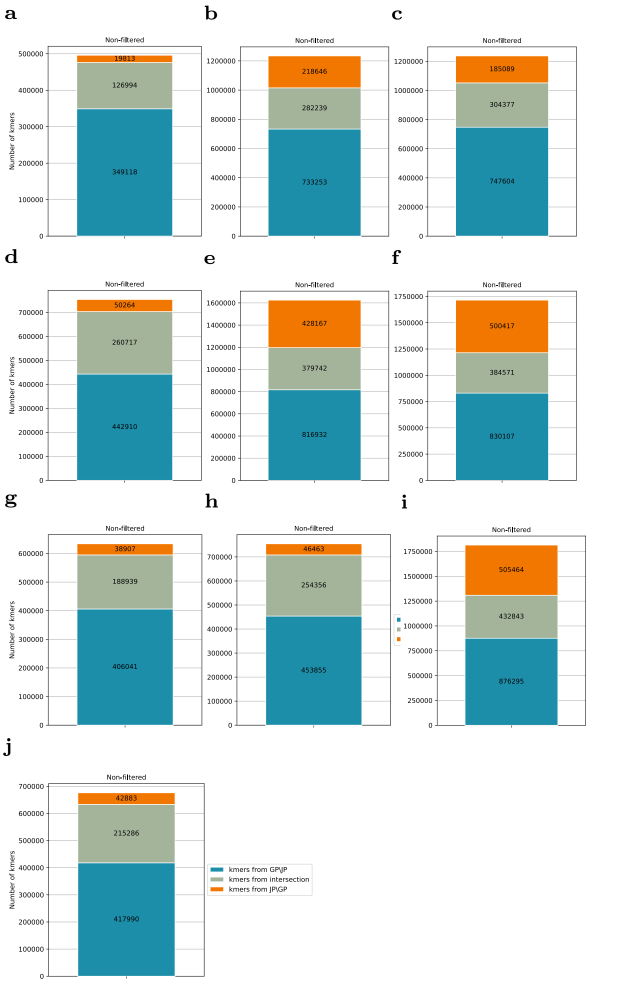


**Fig. 1. Number of 9-mers after the generation step and before the GTEx filtering.** (A-J) Samples TCGA-AO-A0JM, TCGA-24-1431, TCGA-25-1313, TCGA-BH-A18V (BRCA) and TCGA-25-1319, TCGA-61-2008, TCGA-C8-A12P, TCGA-A2-A0SX, TCGA-24-2298, TCGA-A2-A0D2 (OV)

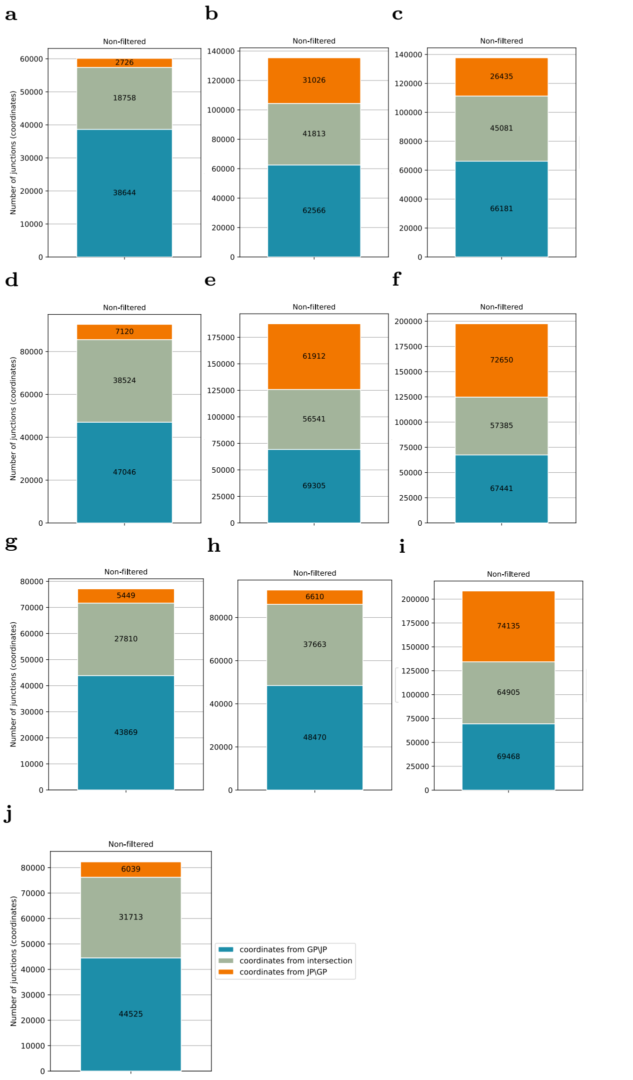


**Fig. 2. Number of junctions after the generation step and before the GTEx filtering.** (A-J) Samples TCGA-AO-A0JM, TCGA-24-1431, TCGA-25-1313, TCGA-BH-A18V (BRCA) and TCGA-25-1319, TCGA-61-2008, TCGA-C8-A12P, TCGA-A2-A0SX, TCGA-24-2298, TCGA-A2-A0D2 (OV)

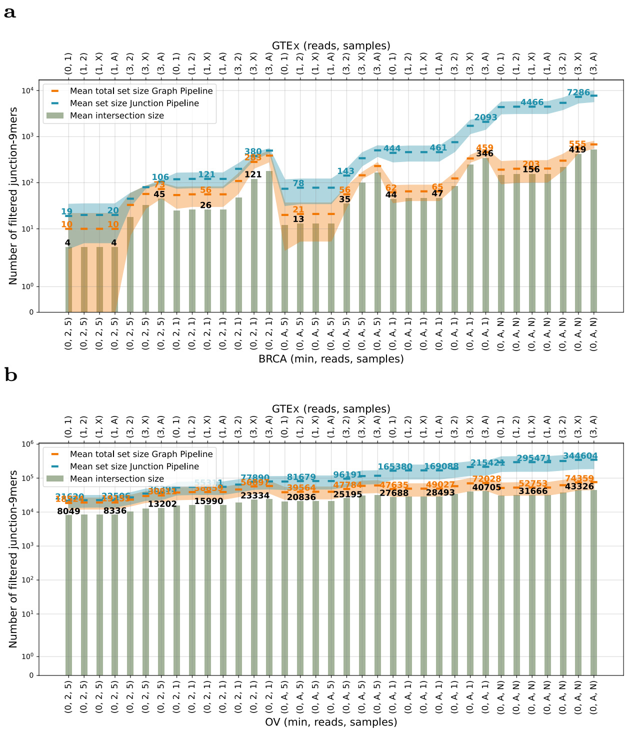


**Fig. 3. Effect of a subset of filtering parameters on the number of 9-mer candidate neoantigens.** Numbers of 9-mer candidates from the GP (orange) and JP (blue), shown in log base 10 scale for BRCA (a) and OV (b) samples. Within each plot, the mean value for one parameter set and one pipeline across samples is shown as a horizontal line within a ribbon representing the range of values across the 5 samples. Sage green bar plots show the mean size of the intersection between the two pipelines for each parameter set. The filter sets shown are ordered from most stringent (left) to most lenient (right). These are ordered first by cancer truth/foreground parameters (listed bottom, where the three comma-separated values are, in order, the numbers of minimum target sample reads required, reads across the cohort samples, and number of cohort samples respectively) and then by cancer specificity/background parameters (listed top, where the two comma-separated values are the number of normal reads allowed and then the number of normal samples allowed). Numerical values under 10 are reported directly; 10 is written as 'X', Any' as 'A' and 'None' as 'N'. More information in Methods Table 1.

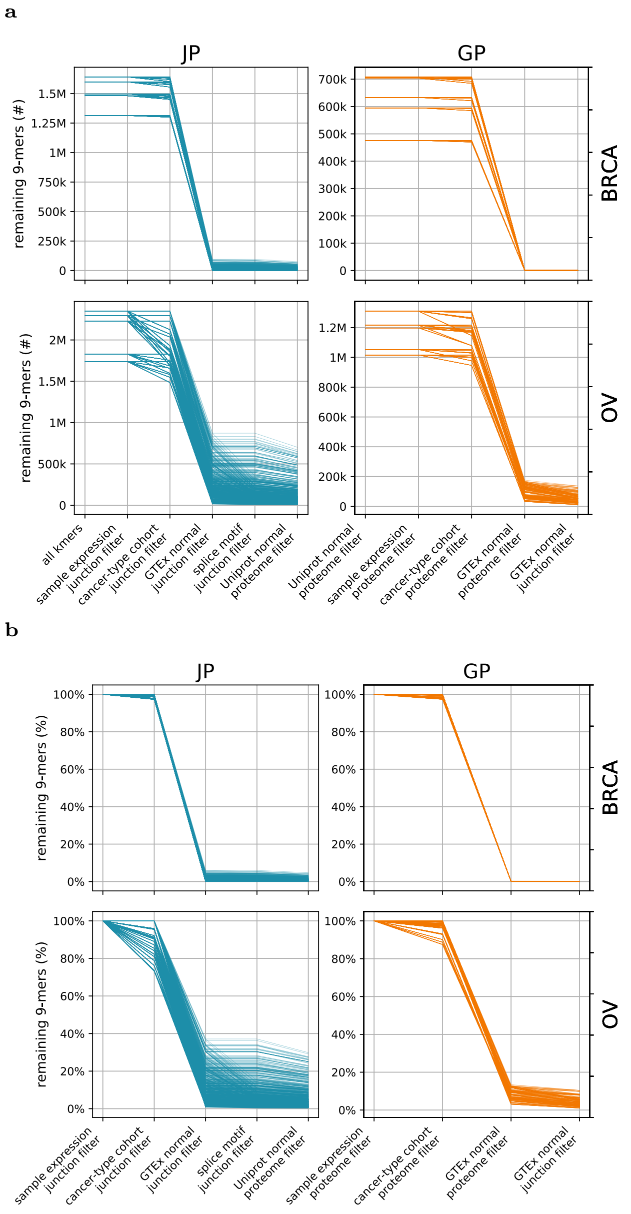


**Fig. 4. Sequential effect of the applied filters on the number of 9-mer candidate neoepitopes.** (a) The absolute number and (b) the percent of the initial -9-mer candidates remaining in each filtering experiment at each sequential filtering step, shown for BRCA samples (top) and OV samples (bottom) for each of the JP (blue, left) and GP (orange, right) pipelines. Within each panel, each line represents the 9-mers present in one filtering experiment performed on one sample from start to finish, with 75 experiments per sample for the GP (750 total) and 180 total experiments for the JP (1800 total). This figure represents the full breadth of experiments run, including canonical splice motif and tissue-matched normal filtering for the JP, and a more broad range of normal expression levels for the GP.

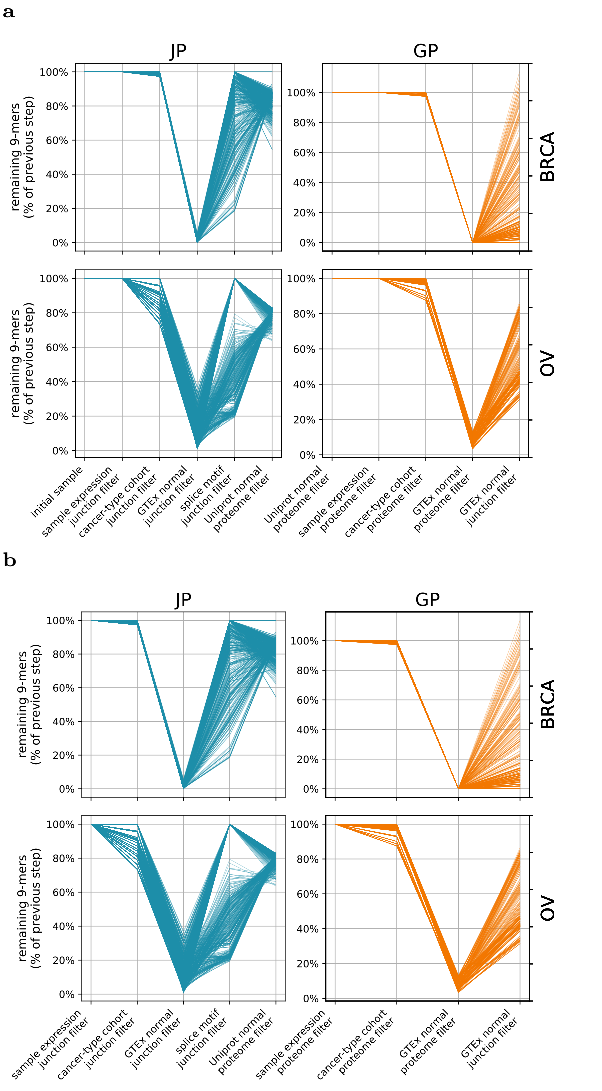


**Fig. 5. Relative sequential effect of the applied filters on the number of 9-mer candidate neoepitopes.** (a) Percent of 9-mer candidates from previous step including the generation step or (b) starting at the filtering step, shown for BRCA samples (top) and OV samples (bottom) for each of the JP (blue, left) and GP (orange, right) pipelines. Within each panel, each line represents the 9-mers present in one filtering experiment performed on one sample from start to finish, with 75 experiments per sample for the GP (750 total) and 180 total experiments for the JP (1800 total). This figure represents the full breadth of experiments run, including canonical splice motif and tissue-matched normal filtering for the JP, and a more broad range of normal expression levels for the GP.

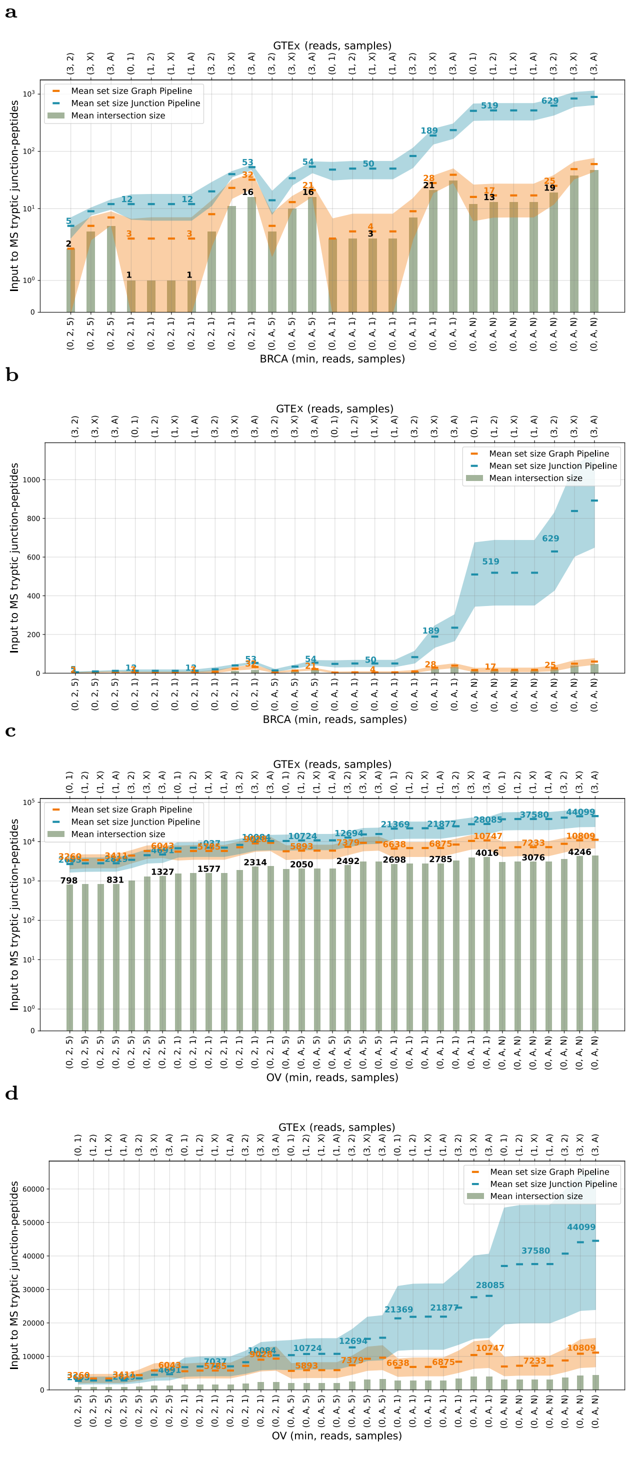


**Fig. 6. Number of tryptic junction peptides provided as input to the proteomics methods.** 2, 3 or 4-exons peptides which contained a candidate neoepitope sequence (9-mers passing GTEx and cancer filters) were selected. The peptides were trypsine digested and the tryptic fragments overlapping the candidate junction were extracted. Log10 of the number of input tryptic junction peptides in BRCA (a) and OV (c) samples. Number of input tryptic junction peptides in BRCA (a) and OV (c) samples. The filter sets shown are ordered from most stringent (left) to most lenient (right). These are ordered first by cancer truth/foreground parameters (listed bottom, where the three comma-separated values are, in order, the numbers of minimum target sample reads required, reads across the cohort samples, and number of cohort samples respectively) and then by cancer specificity/background parameters (listed top, where the two comma-separated values are the number of normal reads allowed and then the number of normal samples allowed). Numerical values under 10 are reported directly; 10 is written as 'X', Any' as 'A' and 'None' as 'N'. More information in Methods Table 1.

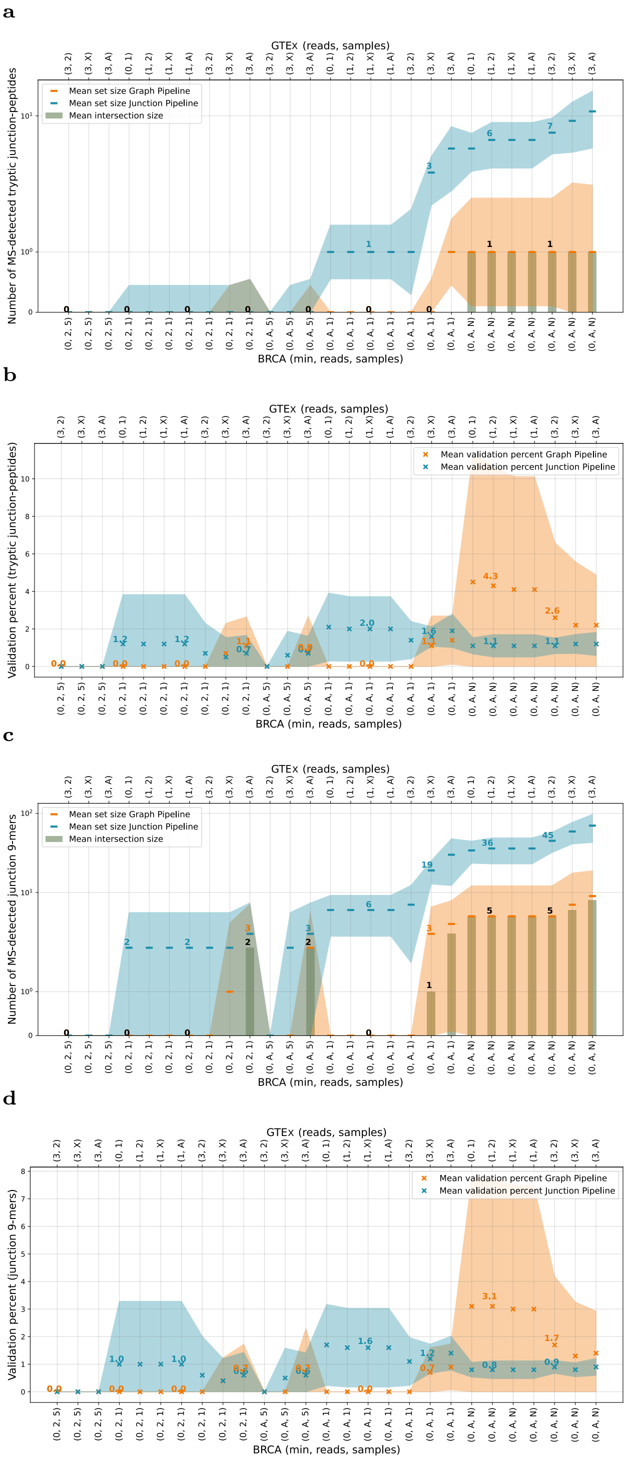


**Fig. 7. Proteomics validation for BRCA samples after *joint* pipeline search.** Candidate sets for both pipelines and experiments were pooled. *Subset-neighbor search* was applied and the peptide-PSM-FDR was calculated with *Crema*. (a) Number of tryptic-junction peptides validated with MS. (b) Validation percent for tryptic-junction peptides. (c) Number of junction-9-mers validated with MS. candidates. (d) Validation percent for junction-9-mers. The filter sets shown are ordered from most stringent (left) to most lenient (right). These are ordered first by cancer truth/foreground parameters (listed bottom, where the three comma-separated values are, in order, the numbers of minimum target sample reads required, reads across the cohort samples, and number of cohort samples respectively) and then by cancer specificity/background parameters (listed top, where the two comma-separated values are the number of normal reads allowed and then the number of normal samples allowed). Numerical values under 10 are reported directly; 10 is written as 'X', Any' as 'A' and 'None' as 'N'. More information in Methods Table 1.

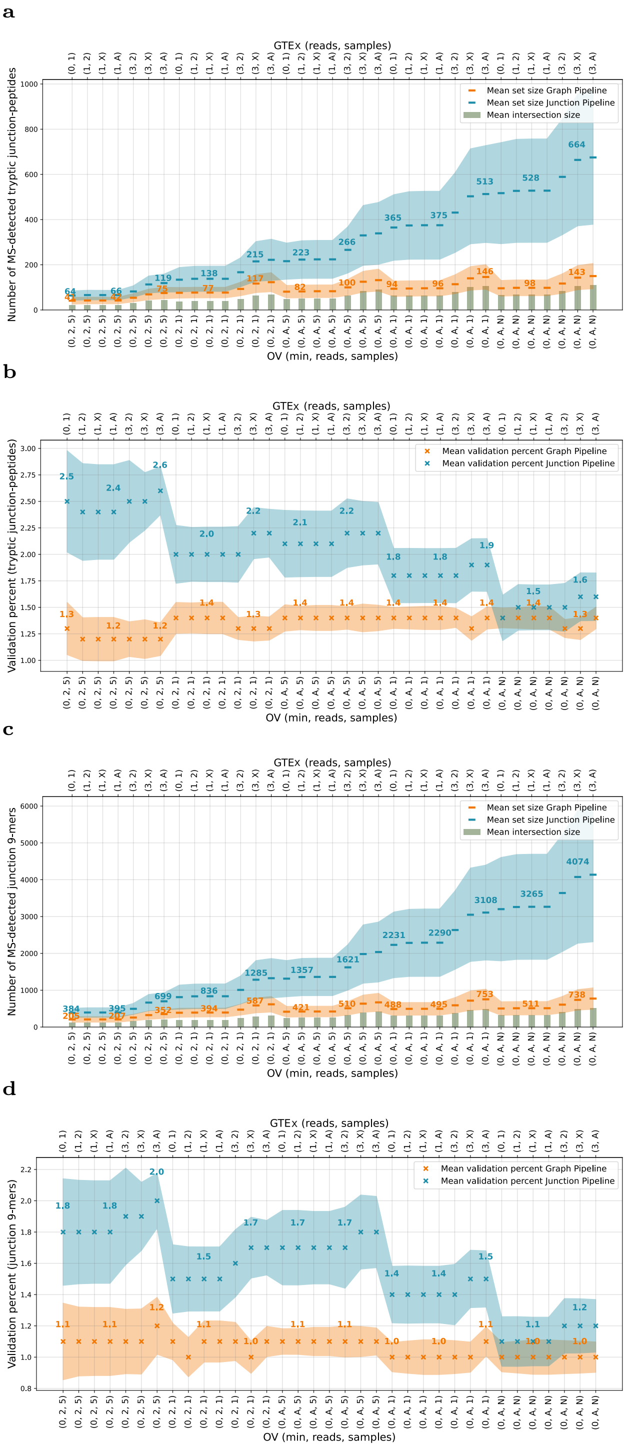


**Fig. 8. Proteomics validation for OV samples after *joint* pipeline search.** Candidate sets for both pipelines and experiments were pooled. *Subset-neighbor search* was applied and the peptide-PSM-FDR was calculated with *Crema*. (a) Number of tryptic-junction peptides validated with MS. (b) Validation percent for tryptic-junction peptides. (c) Number of junction-9-mers validated with MS. candidates. (d) Validation percent for junction-9-mers. The filter sets shown are ordered from most stringent (left) to most lenient (right). These are ordered first by cancer truth/foreground parameters (listed bottom, where the three comma-separated values are, in order, the numbers of minimum target sample reads required, reads across the cohort samples, and number of cohort samples respectively) and then by cancer specificity/background parameters (listed top, where the two comma-separated values are the number of normal reads allowed and then the number of normal samples allowed). Numerical values under 10 are reported directly; 10 is written as 'X', Any' as 'A' and 'None' as 'N'. More information in Methods Table 1.

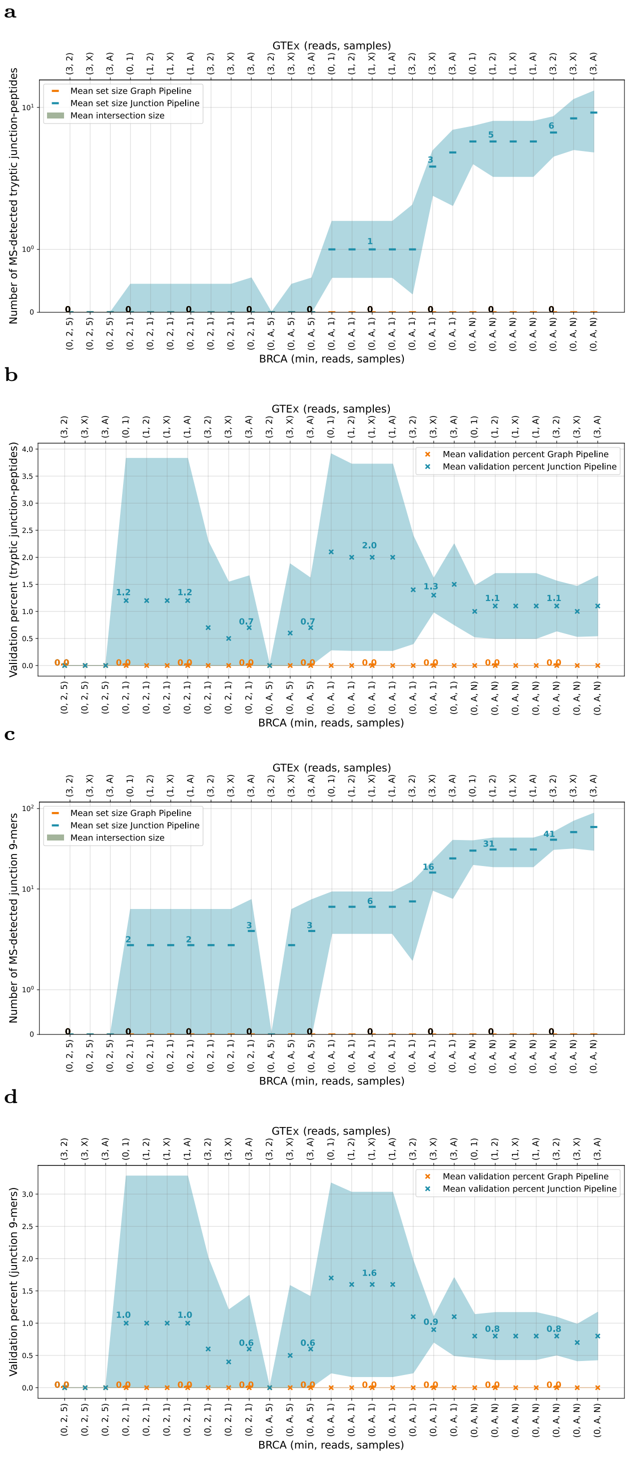


**Fig. 9. Proteomics validation for BRCA samples after *separate* pipeline search.** The pipelines where searched separately while experiments were pooled. *Subset-neighbor search* was applied and the peptide-PSM-FDR was calculated with *Crema*. (a) Number of tryptic-junction peptides validated with MS. (b) Validation percent for tryptic-junction peptides. (c) Number of junction-9-mers validated with MS. candidates. (d) Validation percent for junction-9-mers. The filter sets shown are ordered from most stringent (left) to most lenient (right). These are ordered first by cancer truth/foreground parameters (listed bottom, where the three comma-separated values are, in order, the numbers of minimum target sample reads required, reads across the cohort samples, and number of cohort samples respectively) and then by cancer specificity/background parameters (listed top, where the two comma-separated values are the number of normal reads allowed and then the number of normal samples allowed). Numerical values under 10 are reported directly; 10 is written as 'X', Any' as 'A' and 'None' as 'N'. More information in Methods Table 1.

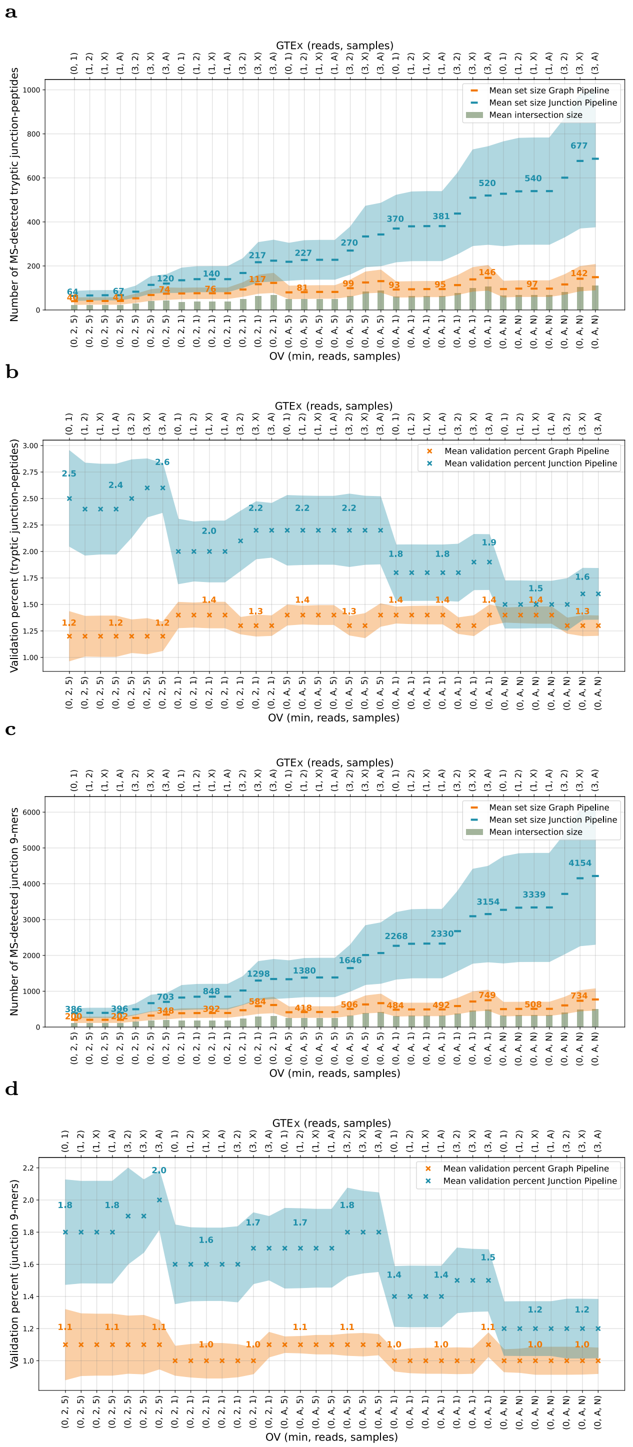


**Fig. 10. Proteomics validation for OV samples after *separate* pipeline search.** The pipelines where searched separately while experiments were pooled. *Subset-neighbor search* was applied and the peptide-PSM-FDR was calculated with *Crema*. (a) Number of tryptic-junction peptides validated with MS. (b) Validation percent for tryptic-junction peptides. (c) Number of junction-9-mers validated with MS. candidates. (d) Validation percent for junction-9-mers. The filter sets shown are ordered from most stringent (left) to most lenient (right). These are ordered first by cancer truth/foreground parameters (listed bottom, where the three comma-separated values are, in order, the numbers of minimum target sample reads required, reads across the cohort samples, and number of cohort samples respectively) and then by cancer specificity/background parameters (listed top, where the two comma-separated values are the number of normal reads allowed and then the number of normal samples allowed). Numerical values under 10 are reported directly; 10 is written as 'X', Any' as 'A' and 'None' as 'N'. More information in Methods Table 1.
